# Plastomes of *Garcinia mangostana* L. and comparative analysis with other *Garcinia* species

**DOI:** 10.1101/2022.02.22.481552

**Authors:** Ching-Ching Wee, Nor Azlan Nor Muhammad, Vijay Kumar Subbiah, Masanori Arita, Yasukazu Nakamura, Hoe-Han Goh

## Abstract

The two varieties of mangosteen (*Garcinia mangostana* L.) cultivated in Malaysia are known as Manggis and Mesta. The latter is preferred for its flavor, texture, and seedlessness. Here, we report a complete plastome (156,580 bp) of the Mesta variety which was obtained through a hybrid assembly approach using PacBio and Illumina sequencing reads. It encompasses a large single-copy (LSC) region (85,383 bp) and a small single-copy (SSC) region (17,137 bp) that are separated by 27,230 bp of inverted repeat (IR) regions at both ends. The plastome comprises 128 genes, namely 83 protein-coding genes, 37 tRNA genes, and 8 rRNA genes. The plastome of the Manggis variety (156,582 bp) obtained from reference-guided assembly of Illumina reads was found to be nearly identical to Mesta except for two indels and the presence of a single nucleotide polymorphism (SNP). Comparative analyses with other publicly available *Garcinia* plastomes, including *G. anomala*, *G. gummi-gutta*, *G. mangostana* var. Thailand, *G. oblongifolia*, *G. paucinervis*, and *G. pedunculata* found that the gene content, gene order, and gene orientation were highly conserved among the *Garcinia* species. Phylogenomic analysis divided the six *Garcinia* plastomes into three groups with the Mesta and Manggis varieties clustered closer to *G. anomala*, *G. gummi-gutta*, and *G. oblongifolia*, while the Thailand variety clustered with *G. pedunculata* in another group. These findings serve as future references for the identification of species or varieties and facilitate phylogenomic analysis of lineages from the *Garcinia* genus to better understand their evolutionary history.

## Introduction

Mangosteen (*Garcinia mangostana* L.) is well-known as the ‘queen of fruits’ and is mainly found in Southeast Asia, particularly in Malaysia, Indonesia, and Thailand^1^. It is priced for its unique taste and valuable natural compounds. Xanthones, which are abundantly found in the ripe fruit pericarp, have been shown to possess antioxidant, anti-cancer, anti-inflammatory, anti-bacterial, and anti-viral properties^2^.

The geographical origin of *G. mangostana* is still under debate. Unlike other flowering plants, *G. mangostana* reproduces apomictically by adventitious embryony in the mother plant without fertilization^3^ and produces *Garcinia-type* recalcitrant seeds without embryo^4^. Morphological and phylogenetic analyses have been performed to examine the parental origin of *G. mangostana* and its relationship with other *Garcinia* species based on internal transcribed spacer (*ITS*)^5–7^, granule-bound starch synthase (*GBSSI*)^7^, *trnS-trnG*, and combination of *trnS-trnG* with *trnD-trnT*^8^. They showed that *G. mangostana* was closely related to *G. malaccensis* and as such, were postulated to have been derived from the hybridization of *G. hombroniana* and *G. malaccensis*^9^. However, as there was only one mangosteen sample (*G. mangostana* TH3) that showed heterozygosity in the *ITS* gene, Nazre proposed *G. mangostana* and *G. malaccensis* to be grouped as one species but different varieties^10^. Mangosteen was suggested to have originated either from the hybrid of different varieties of *G. malaccensis* or the end product of agricultural selective breeding retaining only superior female plants of *G. malaccensis*^10^.

Nonetheless, several reports showed that molecular markers from the nuclear genome could not provide sufficient information for phylogeny demarcation^11^. This is likely due to recombination events in the plant nuclear genome during reproduction^12^. In contrast, the majority of the plastome is inherited maternally. Hence, plastome with a slower rate of evolution provides a better resolution in examining species phylogenetic relationships, adaptive evolution, and divergence dating^13,14^. Recently, with the advancement of next-generation sequencing and long-read sequencing technology, complete plastomes can be obtained easily at low costs. A complete plastome of *G. mangostana* of an unspecified variety that originated from Thailand was first reported in 2017 with the accession number KX822787^15^ (herein, denoted as the Thailand variety) and was shown to be closely related to *G. pedunculata^15^*. In Malaysia, two varieties of *G. mangostana* (Manggis and Mesta) were sequenced and deposited to GenBank^16–18^. However, a complete analysis of the plastomes from these two varieties is yet to be reported.

In this study, we assembled and analyzed the complete mangosteen plastomes of Manggis and Mesta. Furthermore, we performed a comprehensive comparison of all available *Garcinia* plastomes from GenBank and provided an update to a previous comparative analysis^19^. The current comparative study elucidates the structural differences in plastomes for evolutionary inference of the *Garcinia* genus.

## Results

### Characterization of the Mesta plastome

*De novo* assembly of PacBio subreads data using CANU assembler and error-correction with Illumina data using Pilon program produced a total of 7,616 contigs with an N50 genome length of 10,212 bp (Table S1). There was one contig (tig00037541_pilon) with the size of ~165 kb that showed high similarity with the reference plastome in BLAT analysis. It was a circular contig as indicated by the dot plot analysis using Gepard^20^ (Figure S1). One of the identical overlapping ends (~9.3 kb) was trimmed and 18 bases were manually added based on Illumina read correction to obtain the final Mesta plastome size of 156,580 bp. The average coverage of Mesta plastome with PacBio subreads and Illumina clean reads was 265X and 3,751X, respectively (Figure S2 & Table S2).

The Mesta plastome constituted a typical conserved quadripartite structure with two inverted repeat (IR) regions (each 27,030 bp) separating the large single-copy (LSC) region (85,383 bp) from the small single-copy (SSC) region (17,137 bp) (Figure 1). The average GC content of the plastome was 36.2% while, the GC content in LSC, SSC, and IR regions was 33.6%, 30.2%, and 42.2%, respectively. A total of 128 genes were identified, including 77 unique protein-coding genes with 6 duplicated genes in IR; 30 unique tRNAs (7 duplicated genes in IR), and 4 rRNAs (4 duplicated genes in IR) (Table 1 & Table 2).

**Figure 1.**
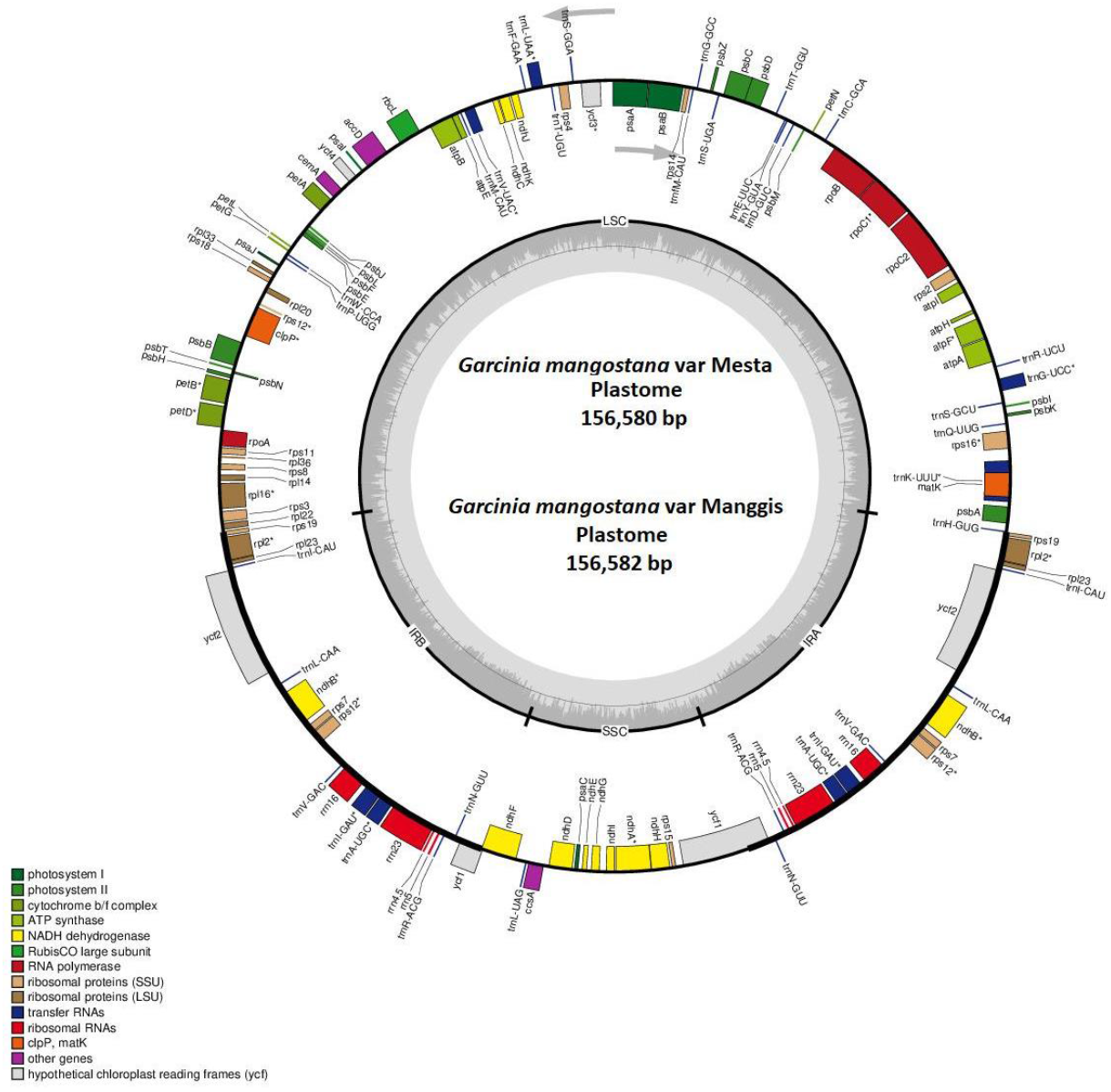
The circular plastome of *G. mangostana* variety Mesta and Manggis. Genes inside the circle are transcribed clockwise while genes outside the circle are transcribed anti-clockwise as indicated by the gray arrows. The gray bars inside the circle represent the GC content of the sequence. Asterisks (<) indicate genes containing intron(s).

### Manggis plastome assembly

The Manggis Illumina clean reads had a higher mapping rate to the Mesta plastome than the Thailand variety (Figure S3 & Table S3). Hence, the Mesta plastome was used for a reference-guided genome assembly of the Manggis plastome. The complete plastome of Manggis had a genome size of 156,582 bp (Table 1) after manual curation with the same genome features as observed in the Mesta plastome (Figure 1) except for one single nucleotide polymorphism (SNP) and two indels (Figure S4 & Table S4). The average coverage of the Manggis plastome with the Illumina short reads was 2,636X (Table S3).

**Table 1.** Summary statistics of plastomes from different *Garcinia* species.

| Species |  | Plastome size (bp) | Size (bp) |  |  | Number of genes* |  |  | GC content (%) |  |  |  |  |
| --- | --- | --- | --- | --- | --- | --- | --- | --- | --- | --- | --- | --- | --- |
|  |  |  | LSC | SSC | IR | All | Protein-coding | rRNA | tRNA | All | LSC | SSC | IR |
| *G. anomala* |  | 156,774 | 85,586 | 17,082 | 27,053 | 128 (111) | 83 (77) | 8 (4) | 37 (30) | 36.1 | 33.5 | 30.3 | 42.1 |
| *G. gummi-gutta* |  | 156,202 | 84,996 | 17,088 | 27,059 | 127 (110) | 83 (77) | 8 (4) | 36 (29) | 36.2 | 33.5 | 30.3 | 42.1 |
| *G. mangostana* | Manggis | 156,582 | 85,385 | 17,137 | 27,030 | 128 (111) | 83 (77) | 8 (4) | 37 (30) | 36.2 | 33.6 | 30.2 | 42.2 |
|  | Mesta | 156,580 | 85,383 | 17,137 | 27,030 | 128 (111) | 83 (77) | 8 (4) | 37 (30) | 36.2 | 33.6 | 30.2 | 42.2 |
|  | Thailand | 158,179 | 86,458 | 17,703 | 27,009 | 128 (111) | 83 (77) | 8 (4) | 37 (30) | 36.1 | 33.5 | 30.1 | 42.2 |
| *G. oblongifolia* |  | 156,577 | 85,393 | 17,064 | 27,060 | 128 (111) | 83 (77) | 8 (4) | 37 (30) | 36.2 | 33.6 | 30.3 | 42.2 |
| *G. paucinervis* |  | 157,702 | 85,989 | 17,737 | 26,988 | 128 (111) | 83 (77) | 8 (4) | 37 (30) | 36.2 | 33.6 | 30.3 | 42.2 |
| *G. pedunculata* |  | 157,688 | 85,998 | 17,656 | 27,017 | 128 (111) | 83 (77) | 8 (4) | 37 (30) | 36.2 | 33.6 | 30.2 | 42.2 |

\*Parentheses indicate the number of unique genes

**Table 2.**
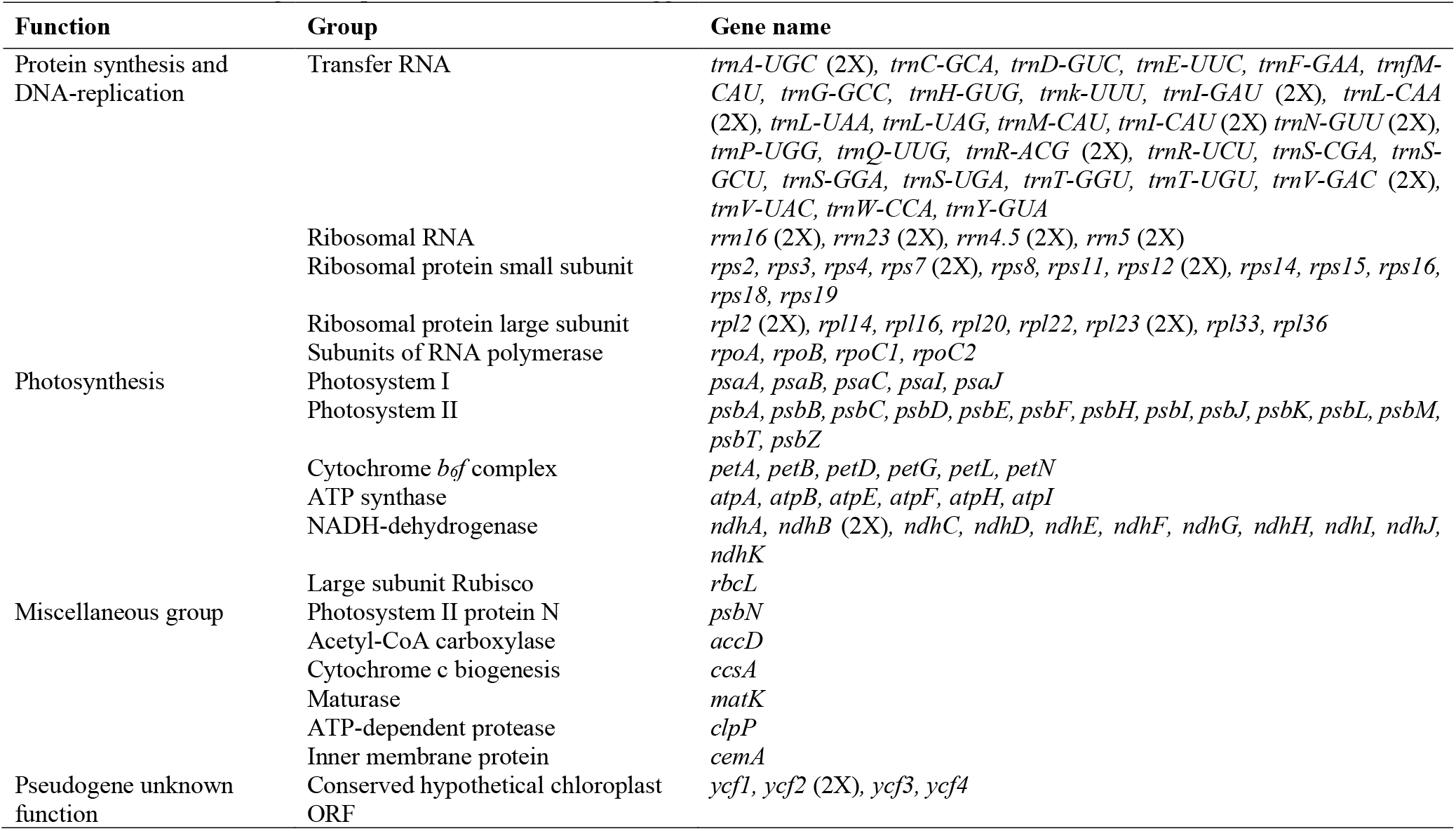
List of annotated genes in plastomes of Mesta and Manggis.

### Plastome feature comparison

In comparison with the plastomes of other *Garcinia* species, the plastome sizes of both Mesta and Manggis varieties (156,580 bp and 156,582 bp, respectively) were larger than *G. gummi-gutta* (156,202 bp) and *G. oblongifolia* (156,577 bp), but smaller than *G. anomala* (156,774 bp), *G. pedunculata* (157,688 bp), *G. paucinervis* (157,702 bp), and the Thailand variety (158,179 bp).

Gene *infA* was found in GenBank for *G. pedunculata* (NC_048983) and *G. anomala* (MW582313), while gene *rpl32* was found for *G. pedunculata* (NC_048983). However, both genes were not annotated accurately. Multiple sequence alignment of the *infA* and *rpl32* genes (Figure S5) with other species showed that both annotated sequences did not have a conserved region as compared with other species. Hence, these genes were re-annotated and revised for accuracy prior to subsequent comparative analysis.

The number of protein-coding genes (83 CDS) was the same for all the *Garcinia* species. However, gene *trnH-GUG* was not found in *G. gummi-gutta*. Hence, the total number of genes for *G. gummi-gutta* was 127 compared to 128 genes for other *Garcinia* species (Table 1). The overall GC content (36.1-36.2%) and GC content found in LSC (33.5-33.6%), SSC (30.1-30.3%), and IR (42.1-42.2%) were similar among the *Garcinia* species.

For *Garcinia* plastomes, there were a total of 18 unique genes (*rps16, atpF, rpoC1, ycf3, rps12, clpP, petB, petD, rpl16, rpl2, ndhB, ndhA, trnK-UUU, trnG-UCC, trnL-UAA, trnV-UAC, trnI-GAU*, and *trnA-UGC*) containing at least one intron with two introns in *clpP* and *rps12* (Table S5), which is similar to those generally found in other plants^21^. Gene *clpP* was located in the LSC region. Meanwhile, the 5’ exon of the *rps12* gene was located in the LSC region while the 3’ exon was located in the IR regions, which is commonly observed in plastomes of other species such as, *Rhodomyrtus tomentosa, Salvia* spp^22,23^, *Ananas comosus* var. comosus^21^, and ferns^24^. Among the genes, *trnK-UUU* had the longest intron length, which is in agreement with previous studies^25–27^.

### Codon usage and amino acid frequency

The total number of codons in 83 protein-coding genes found in the plastomes were different among the *Garcinia* species ranging from 26,195 in *G. pedunculata*, 26,216 in Thailand variety, 26,244 in *G. paucinervies*, 26,249 in *G. anomala*, 26,257 in both Manggis and Mesta varieties, 26,265 in *G. gummi-gutta*, to 26,268 in *G. oblongifolia* (Table S6). There were several common findings in the codon usage analysis of *Garcinia* plastomes: (1) a total of 20 translated amino acids; (2) the most frequent amino acid was leucine while the least frequent was cysteine (Figure S6); (3) there were 30 types of codon out of 64 codons with relative synonymous codon usage (RSCU) values >1.0 (ending with either A or U, except for UUG) and 32 types of codons with RSCU < 1.0 (ending with either C or G, except for AUA, CUA, and UGA); (4) both AUG and UGG had an RSCU value = 1 (Table S6). Similar findings were also detected in other plants, such as *Euphorbiaceae* and *Rhodomyrtus tomentosa*^23,28^,

Generally, the start codon is ATG but there were exceptions for several genes with initiation codons of GTG or ACG due to the RNA editing events during transcription. The first discovery of such an event came from the study of *rpl2* gene in the maize plastome when the start codon of this gene changed from ACG to ATG during transcription^29^. Other examples include GTG as an initiation codon of *psbC* gene^30^ and ACG for *ndhD* gene^31^ in the tobacco plastome, start codons ACG and GTG for *rpl2* and *rps19* genes, respectively in *Oryza sativa* plastome^32^. Similarly, there were three genes (*rpl33, rps19*, and *ndhD*) in the plastomes of *Garcinia* species that did not start with ATG. The start codon was GTG for both *rpl33* and *rps19* genes in all the *Garcinia* species, except for *rps19* in *G. anomala* which starts with an ATG. As for *ndhD*, ATG was found in *G. anomala* while TTG was found in *G. pedunculata* and ACG for all the other *Garcinia* species.

### Simple Sequence Repeat (SSR) analysis

A total of 88 SSRs were identified, including 79 mononucleotide repeats, 7 dinucleotide repeats, and two trinucleotide repeats with a total sequence size of 1,065 bp and 1,067 bp for Mesta and Manggis varieties, respectively. The SSRs identified in the plastomes of both Mesta and Manggis were the same except Manggis had additional mononucleotide A (SSR no 3 at Table S7 & S8) and C (SSR no 25 at Table S7 & S8), respectively, compared to Mesta. There were 14 SSRs with less than 100 bp in length between two SSRs (compound microsatellite) in both Mesta and Manggis plastomes. The most abundant motif found in both Mesta and Manggis varieties was mononucleotide repeats (89.8%), in which mononucleotide T (48.9%) and A (38.6%) constituted the highest portion compared to mononucleotide C (1.1%) and G (1.1%). Among them, only 10 of the mononucleotide repeats were located at the coding regions of *rpoC2, rpoC1, rpoB, rps19, ycf2*, and *ycf1* genes. All dinucleotide and trinucleotide repeats were located at the non-coding regions, which are common phenomena in flowering plants^33^. All the SSRs in the plastomes of Mesta and Manggis varieties including their respective locations are listed in Table S7 and Table S8, respectively.

The total number of SSR varied among the different *Garcinia* species (Figure 2). Mesta and Manggis varieties had the lowest number of total SSR (88) compared to *G. oblongifolia* (106), Thailand variety (105)*, G. gummi-gutta* (102), *G. anomala* (96), *G. paucinervis* (94), and *G. pedunculata* (91). Mononucleotide repeats constituted the highest percentage in SSR analysis in this study, which is in agreement with previous studies of plastomes from 164 lower and higher plants^33^. The highest dinucleotide repeat was found in *G. oblongifolia* (11), followed by Thailand variety (8) and Mesta/Manggis varieties (7) while the other *Garcinia* species showed 6 dinucleotide repeats. The Thailand variety had 3 trinucleotide repeats, which was the highest among all the *Garcinia* species. There was only one trinucleotide repeat found in both *G. gummi-gutta* and *G. paucinervis* compared to two trinucleotide repeats found in Mesta, Manggis, and *G. oblongifolia*. Trinucleotide repeat was not found in *G. anomala* and *G. pedunculata*. The majority of the mononucleotide SSR belonged to the A/T repeats and the same findings were observed in the previous studies^21,23,26^.

**Figure 2.**
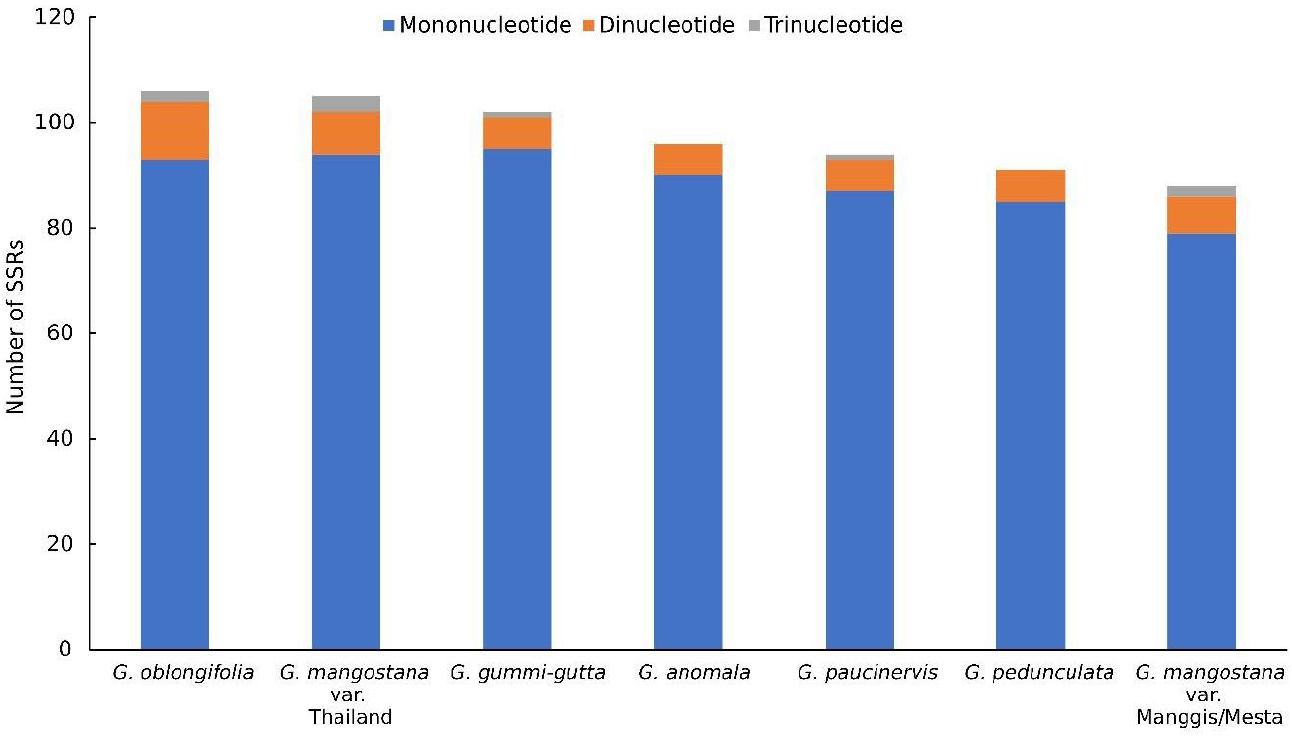
SSR analysis of six *Garcinia* species plastomes.

### Long Repeat Analysis

By using the default setting of 50 for maximum computed repeats, different types of long repeats were detected in the plastomes of *Garcinia* species, including forward repeats, reverse repeats, complement repeats, and palindromic repeats (Figure 3). The palindromic repeat was the most common repeat found in *Garcinia* species followed by the forward repeats, which was also observed in other plants^27^. Mesta and Manggis varieties had the highest number of palindromic repeats of 31, while *G. anomala* had the lowest number of palindromic repeats of 25. The reverse repeat was found in all *Garcinia* species, except *G. gummi-gutta* which had the highest number of forward repeats (21). *G. paucinervis* had the highest number of reverse repeats (5), followed by three reverse repeats found in the Thailand variety and *G. anomala* while the other *Garcinia* species only had one reverse repeat. In addition, complement repeat was only found in *G. anomala* (2), Mesta and Manggis varieties (1), and *G. paucinervis* (1).

**Figure 3.**
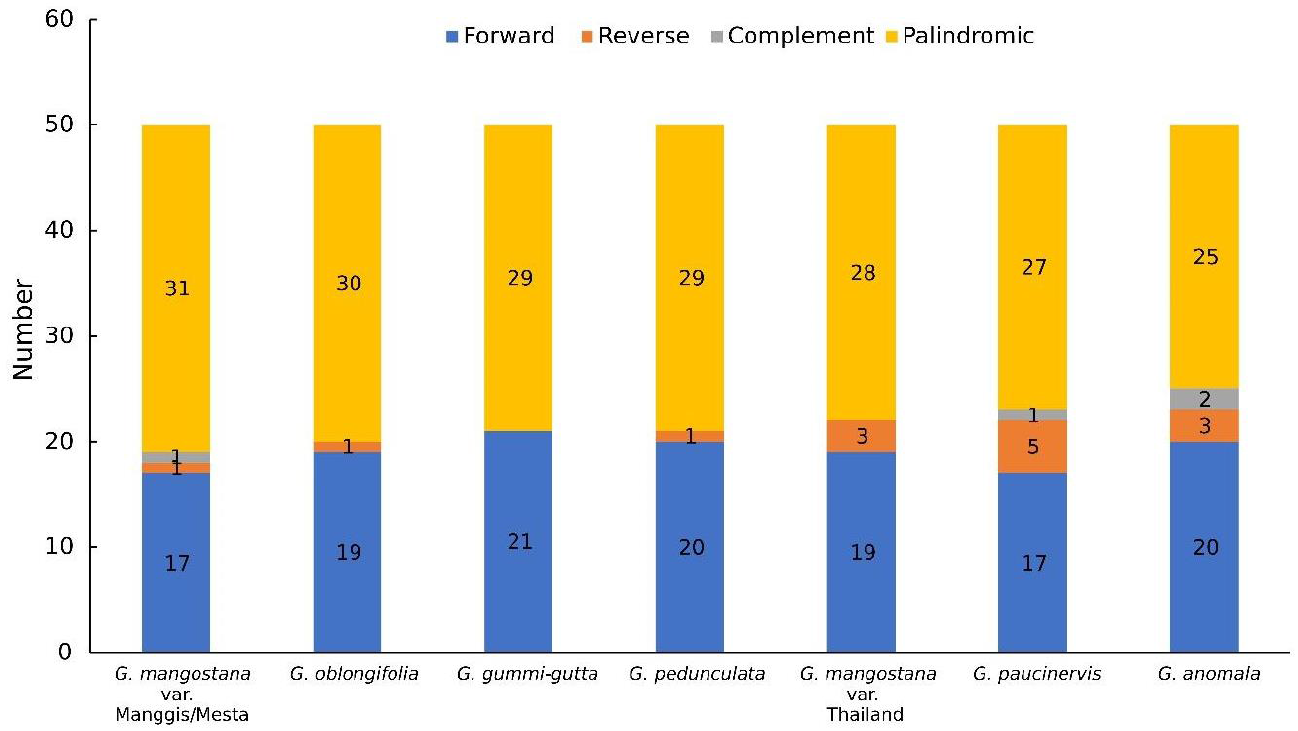
Long repeat analysis of six *Garcinia* species plastomes.

### Contraction and Expansion of the Inverted Repeat Region

The smallest inverted repeat (IR) was found in *G. paucinervis* (26,988 bp) while the largest IR was found in *G. oblongifolia* (27,060 bp) (Figure 4). Genes that can be found at or close to the junctions of IRs were *rps19, ndhF, ycf1*, and *trnH*. Gene *rps19* was located at the LSC/IRb junction site (JS) and the fragment size located at LSC in all *Garcinia* species was 60 bp, except for the Thailand variety which was only 8 bp. In addition, the *rps19* gene fragment of the Thailand variety located at IRb site was 1 bp longer (220 bp) than the *rps19* gene fragment (219 bp) of the other *Garcinia* species at the same location. The *ndhF* gene spanned across the SSC/IRb with 1 bp located at IRb region for *G. anomala, G. gummi-gutta*, Manggis and Mesta varieties, and *G. oblongifolia*. However, it was 1 bp away from the SSC/IRb junction site of both *G. paucinervis* and *G. pedunculata*. Interestingly, Thailand mangosteen was the only one with *ndhF* gene 4 bp from the SSC/IRb junction. The *ycf1* gene fragment (1,421 bp) in the IRa region was the same for all *Garcinia* species, except for *G. pedunculata* and *G. paucinervis* (1,419 bp). The size of the *ycf1* fragment in the SSC region range from 4,204 bp to 4,245 bp. In addition, it was found that *tRNA-trnH* was missing at the IRa/LSC junction of *G. gummi-gutta*. In comparison with *Erythroxylum novogranatense* (Plastome size: 163,937 bp; LSC: 91,383 bp; SSC: 18,138 bp; IR: 27,208 bp), a sister group of *G. mangostana*^15^, plastome size, LSC, SSC, and IR regions of *Garcinia* species were much shorter (Figure 4 & Table 1).

**Figure 4.**
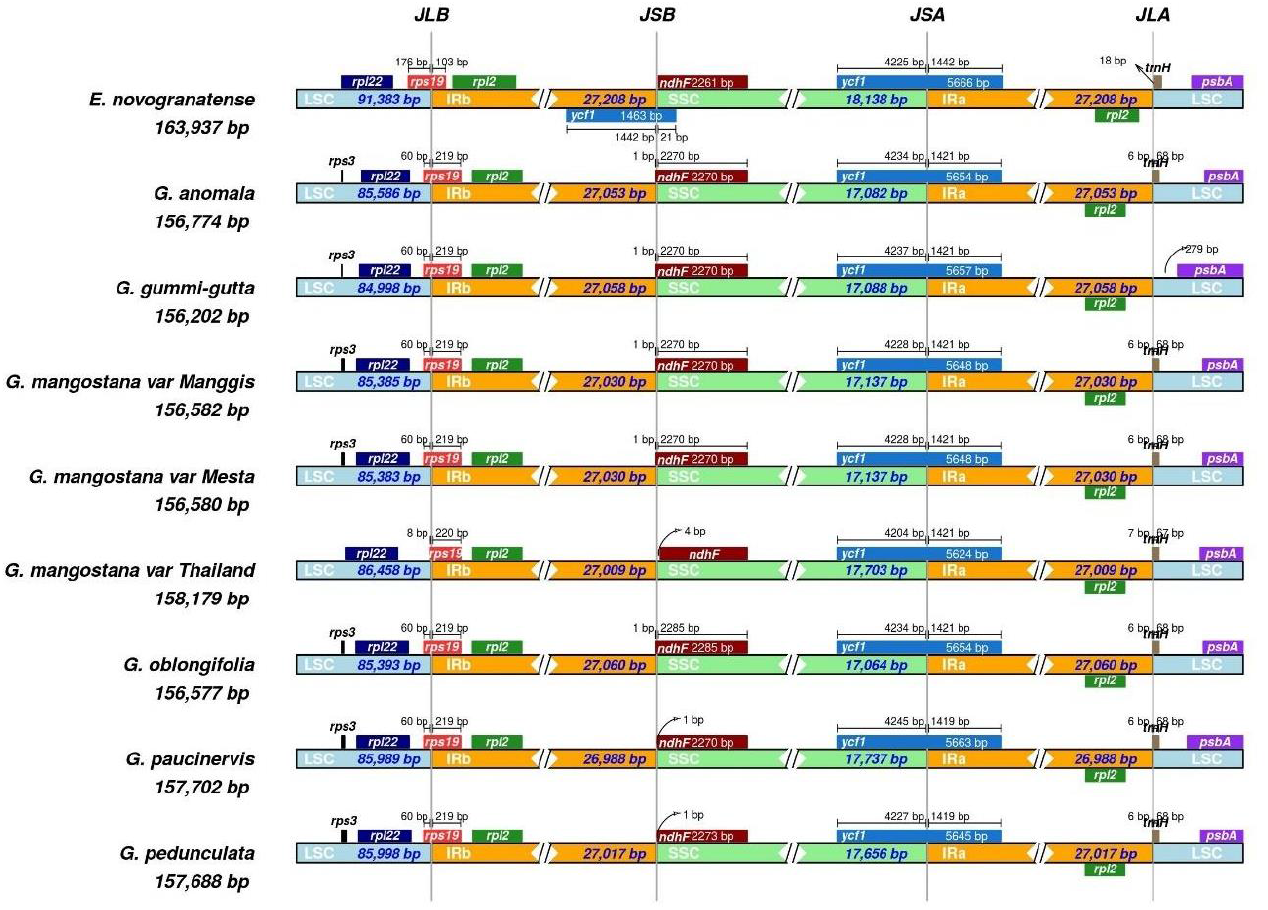
Comparison of genes on the borders of the LSC, SSC, and IR regions among six *Garcinia* plastomes and *Erythroxylum novogranatense*. Corresponding plastome size is shown on the left of each track. The intervals show the distance between the start and end coordinates of a particular gene from the junction sites, namely JLB (LSC/IRb), JSB (SSC/IRb), JSA (SSC/IRa), and JLA (LSC/IRa). The sequence length in each region is annotated for genes spanning the junction sites.

### Comparative Plastome Analysis

Plastomes comparison using Mesta as a reference was performed using mVista online alignment tool (Figure 5). The comparison among *Garcinia* species showed that (1) IR regions were more conserved compared with LSC and SSC regions, (2) coding regions were more conserved than intergenic regions. This result agreed with previous reports in Plantaginaceae, Rosaceae, and Sapindaceae families^25,34,35^. For LSC, highly divergent intergenic regions include *trnH-psbA*, *trnQ-psbK*, *trnG-trnR*, *atpF-atpH*, *atpH-atpI*, *trnT-psbD*, *ndhC-trnV*, *rbcL-accD,psbB-psbT*, and intergenic region within the *rpl16* gene. As for SSC, the highly divergent regions include *ndhF-trnL* and intergenic region within the *ndh*A gene. Divergent regions were also found in the coding regions such as *matK, rpoC2, rpoC1, rpoB, rbcL, accD, ycf4, cernA, petA, petD, rpoA, ndhF, ccsA, ycf1*, and *ycf2* (Figure 5). In comparison with *Erythroxylum novogranatense*, plastome sequences of *Garcinia* species were conserved within the Clusiaceae family.

**Figure 5.**
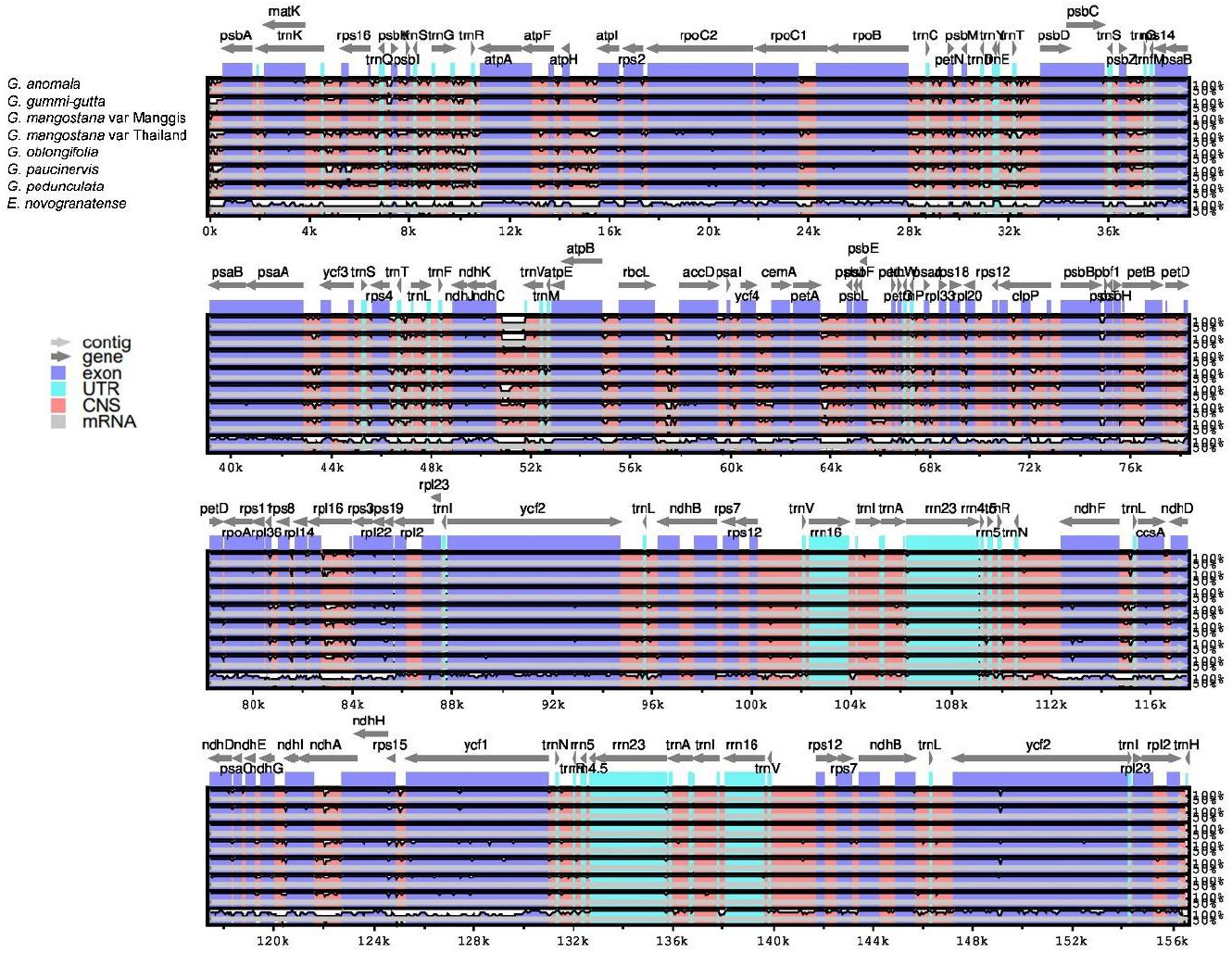
Alignment visualization of *Garcinia* species using Mesta as a reference by using mVista alignment program. CNS: Conserved Non-Coding Sequences; UTR: Untranslated Region. The gray arrows above the alignment indicate the direction of the gene translation. The identity percentage (50-100%) was indicated at the right-side of the mVista plot.

### Phylogenomic Analysis

A total of 74 protein-coding genes (Table S9) were used for phylogenomic analysis. Phylogenomic analysis showed that both Manggis and Mesta varieties were grouped together as the CDS sequences were 100% identical despite some base differences in non-coding regions. Both were grouped under the clade of *Garcinia* species in Malpighiales order among the three groups of *Garcinia* species (Figure 6). *G. anomala* and *G. gummi-gutta* were closely related and formed one group with the Mesta/Manggis varieties and *G. oblongifolia*. The Thailand variety and *G. pedunculata* formed another group, while *G. paucinervis* formed the third group in the Clusiaceae family.

**Figure 6.**
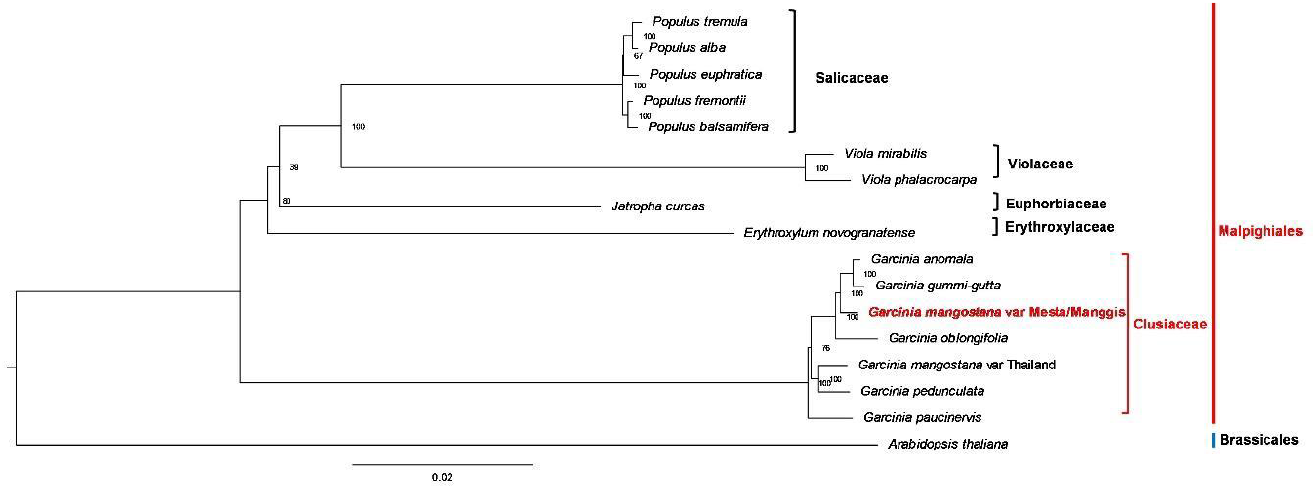
Phylogenetic tree (maximum likelihood) construction of 16 species (three varieties from *G. mangostana*) based on 74 protein-coding genes. The red outer line indicates order name while the inner line indicates the family name.

## Discussion

Plant DNA is rich in plastome (~5-20%) and hence, enrichment strategy is not required for sequencing^14^. In this study, a complete mangosteen plastome of Mesta variety (156,580 bp) was successfully obtained from PacBio long reads. Here, the use of long reads for the assembly of the plastid genomes was ideal to obtain longer contigs and to resolve repetitive regions^26,36,37^. In addition, Illumina sequencing data were used to correct the random errors within PacBio reads^38^. Furthermore, the polished Mesta plastome allowed for the reference-guided assembly of the plastome from Manggis which had only Illumina short reads.

Both the Mesta and Manggis plastomes were nearly identical (Fig. 1), consisting of a typical conserved quadripartite plastome structure found in most of the land plants^14,39^. Generally, the number of genes encoded in a plastome ranges from 110 to 130 genes^35^. Both Malaysian mangosteen plastomes fall within the range with 128 genes (111 unique genes), consistent with all other *Garcinia* species found in GenBank, except for *G. gummi-gutta* which lost one *trnH* gene (Table 1). Typically, plastomes have 30-31 tRNAs^40,41^ and the loss of tRNA genes is not uncommon. For instance, two hemiparasitic *Taxillus* species lost 7 tRNAs, including tRNA-*trnH*^42^. Sometimes, missing tRNAs might be replaced by other types of anticodons, such as *Neochloris aquatica* (NC_029670.1), *Bracteacoccus giganteas* (NC_028586.1), *Tetradesmus obliquus* (NC_008101.1), *Floydiella terrestris* (NC_014346.1), *Schizomeris leibleinii* (NC_015645.1), and *Oedogonium cardiacum* (NC_011031.1)^43^.

The plastome size, structure, and gene content are highly conserved among *Garcinia* species. There were 18 genes (12 protein-coding genes and 6 tRNAs) containing intron(s) in the plastomes of *Garcinia* species. Although introns are not protein-coding, they play an important role in gene expression by regulating the rate of transcription, nuclear export, and stability of transcripts^44,45^. The loss of introns such as *rpl*2 and *rps16* has been reported in the plant plastomes^46–48^ but, we did not find any evidence of this in the plastomes of *Garcinia* species.

We detected two mis-annotations of genes (*infA* and *rpl32*) in *G. pedunculata*^19^ and one mis-annotation of *infA* in *G. anomala^49^*. The *infA* is usually located between *rpl36* and *rps8*, whereas *rpl32* is usually located between *ndhF* and *trnL-UAG*^50^. Instead, the annotated *infA* was located within *rpoC1* gene (*G. anomala:* MW582313^49^; *G. pedunculata:* NC048983^51^) while, *rpl32* of *G. pedunculata* was located between *rpoB* and *trnC-GCA*. Hence, both genes were actually not found in the *Garcinia* plastomes, in agreement with a previous report that both *infA* and *rpl32* genes were lost in *G. mangostana*^52^. Plastid gene transfers to the nuclear genome (e.g., *accD, infA, rpl22*, and *rpl32*) have been documented in several plants ^50,53,54^. The abundance of *infA* and *rpl32* transcripts in the seed transcriptome^4^ suggests the same scenario for *G. mangostana*.

Genetic variations in *G. mangostana* cultivars have been shown by randomly amplified DNA fingerprinting (RAF) and inter simple sequence repeat (ISSR) molecular markers^55–57^. Plastomes also contain SSR and long repeats^58,59^. SSR, which is a stretch of 1-6 bp small repeats, is found extensively in different regions of plastome such as the intergenic regions, intron regions, and protein-coding regions^23^. In contrast, the long repeats found in the plastomes mostly fall within the intergenic region although, some of them were present in protein-coding genes^60^. Repetitive regions might lead to species variation as they have a higher tendency of recombination, translocation, and insertion/deletion^61^. In this study, SSR and long repeat analyses of plastomes showed variations among *Garcinia* species and varieties of *G. mangostana*. This supports that both molecular markers are useful for species identification and taxonomic studies^62–64^.

One of the main factors that contribute to the plastome size differences is the inverted repeat (IR) expansion and contraction^65,66^. For instance, nine genes were transferred from the SSC to the IR region in *Plantago ovata* resulting in an extremely long IR (37.4 kb)^25^ as compared to IR found in the other plastomes which normally range between 25 kb and 30 kb^40^. In contrast, the loss of IR had been reported in the plastomes of 25 durian varieties recently^37^.

The phylogenomic analysis showed identical protein-coding genes between Mesta and Manggis varieties, which implies the same maternal lineage for both varieties from Malaysia compared to the Thailand variety (Figure 6). This result is congruent with the analysis using the whole plastome sequences of 16 species (Figure S7). The Malaysian (Mesta/Manggis) and Thailand mangosteen varieties did not cluster together in the phylogenomic analysis contrary to the initial hypothesis of this study, which assumed a close phylogenomic relationship of the same species. Analysis of polymorphic sites of the 74 CDS used for phylogenomic tree construction showed a total of 559 variable sites, accounting for 0.85% differences between Manggis/Mesta and Thailand varieties (Table S10). This was inconsistent with the clustering of different *G. mangostana* varieties, including the Thailand varieties in a previous phylogenetic study based on the nuclear *ITS* gene^10^.

To further investigate this discrepancy, we obtained the consensus *ITS* gene sequences of both Mesta (accession number: OK576276) and Manggis (accession number: OK576274) varieties by mapping the respective Illumina filtered reads against the published *ITS* gene sequence (accession number: AF367215). We reconstructed a phylogenetic tree^10^ based on the *ITS* gene sequences of other *Garcinia* species found in GenBank, including *G. celebica, G. gummi-gutta, G. hombroniana, G. oblongifolia, G. paucinervis*, and *G. pedunculata* (Table S11). The *ITS* phylogenetic tree (Figure S8) showed that all the *G. mangostana* varieties were grouped with *G. malaccensis* congruent with the results of the previous study^10^. Meanwhile, Mesta and Manggis were distantly related to *G. gummi-gutta, G. oblongifolia, G. paucinervis*, and *G. pedunculata*. Furthermore, Mesta, Masta, and *G. malaccensis* MY4 were clustered together, away from the Manggis variety. This indicates genetic differences between the two varieties despite near-identical plastomes and supports that both varieties might be originated from *G. malaccensis*.

We found that both Manggis and Mesta varieties showed heterozygosity at certain positions of *ITS* gene (Manggis: position 200; Mesta: position 444, 477, and 527) based on Illumina short reads results, respectively (Figure S9 & S10). Out of ten *G. mangostana* reported in the previous study^10^, only one sample (*G. mangostana* TH3) from Thailand showed heterozygosity. Hence, *G. mangostana* may not have been derived from the hybridization of *G. hombroniana* and *G. malaccensis*^9^. Meanwhile, near-identical Mesta and Manggis plastomes (Figure 1 & 6) indicate the same maternal lineage. This illustrates the different evolutionary inferences from the nuclear genome and plastome analysis as plastids are inherited maternally compared to recombination events in nuclear genomes during reproduction^12^.

As hybridization is a common practice in plant breeding to produce hybrids with desirable traits^67^, the genetic variations and heterozygosity observed in this study could be due to the different germplasms. Different germplasms may hybridize via selective breeding and could have produced different varieties of *G. mangostana* in Malaysia and Thailand^10^. However, this remains highly speculative and requires further investigations of mangosteen from different biogeographical origins as well as plastomes of *G. celebica* (syn. *G. hombroniana*), *G. malaccensis*, *G. penangiana*, and *G. opaca* to ascertain their maternal lineages.

## Conclusion

The complete plastomes of both Mesta and Manggis varieties of *G. mangostana* from Malaysia were successfully assembled and analyzed. PacBio long-read sequencing data helped to resolve the repetitive sequences in Mesta. Subsequently, this allowed reference-guided genome assembly of the Manggis plastome. Notably, the Manggis plastome was almost identical with the Mesta plastome compared to the plastome of Thailand variety. Comparative analysis showed that the gene structure, gene content, gene order, and gene orientation of *Garcinia* plastomes were largely conserved, except for one missing *trnH-GUG* gene in *G. gummi-gutta*. Phylogenomic analysis indicated that the Mesta and Manggis varieties were closer to *G. anomala, G. gummi-gutta*, and *G. oblongifolia*, while the Thailand variety clustered with *G. pedunculata*. Phylogenetic analysis based on the nuclear *ITS* sequences delineated the Mesta and Manggis varieties based on differences in their nuclear genomes. This study suggests different origins of the Mesta/Manggis and Thailand varieties. SSR and long repeats of plastomes identified in this study will provide useful biomarkers for species/variety identification and future lineage study of *Garcinia* genus.

## Materials and methods

### Mesta plastome *de novo* genome assembly

Genome sequences of Mesta variety were obtained from the NCBI SRA database with the accession numbers SRX2718652 to SRX2718659 for PacBio long-read data (9.5 Gb)^17^ and SRX270978 for Illumina short reads (50.2 Gb)^18^. CANU v2.0^68^ was used to perform PacBio raw data correction, trimming, and assembly using default parameters with minor modifications (useGrid=false, genomeSize=6g, batMemory=252, batThreads=32, minInputCoverage=0.15, stopOnLowCoverage=0). The draft genome assembly was polished with Illumina data using Pilon v1.23^68^. Candidate plastome contigs were identified by using BLAT v36.0 alignment tool with the previously reported *G. mangostana* (NC_036341.1) as the query. Identified contig was manually curated based on the read coverage to obtain the final plastome of Mesta for subsequent analysis.

### Manggis plastome assembly

Genome sequences of Manggis variety were obtained from the NCBI SRA database with the accession number SRX1426419 for Illumina reads (51.1 Gb)^16^. Different methods were used for Manggis variety plastome assembly: (1) reference-guided assembly using GetOrganelle v1.7.5^69^, (2) *de novo* assembly using GetOrganelle v1.7.5^69^, and (3) *de novo* assembly using Platanus v1.2.4^70^ (Figure S11). To select the reference for reference-guided genome assembly, Manggis clean reads were aligned against the complete plastomes of Mesta and Thailand^15^ varieties using bwa-mem v0.7.17^71^ and samtools v1.1^72^. Next, the mapping coverage was visualized using weeSAM v1.6. The reference with higher percentage coverage was chosen as the final reference for subsequent analysis. Manual curation was performed on the reference-guided assembled Manggis plastome to obtain the final Manggis plastome (Figure S12 & Table S12).

### Plastome annotation

Plastome annotation was performed online using GeSeq (https://chlorobox.mpimp-golm.mpg.de/geseq.html)^73^. Four *Garcinia* species [*Garcinia gummi-gutta* (NC_047250); *Garcinia mangostana* (NC_036341); *Garcinia oblongifolia* (NC_050384); and *Garcinia pedunculata* (NC_048983)] were used as BLAST-like Alignment Tool (BLAT) references. Respective gene annotations were corrected manually. Lastly, the plastome map was generated using Organellar Genome DRAW (OGDRAW v1.3.1) program with default parameters^74^. The annotated plastomes of both Mesta and Manggis varieties were submitted to NCBI with accession numbers MZ823408 and OK572535, respectively.

### Open Reading Frame (ORF) coordinate adjustment

The length of each gene found in *Garcinia* species was compared. Gene alignment was performed to visualize the differences when dissimilarity in gene length was detected by different annotation software. Next, manual coordinate adjustment was performed to standardize the 5’end and the splicing site of these genes (Figure S13 & Table S13). The adjusted OFR coordinates (Supplementary Data) were used for subsequent analysis.

### Identification of Simple-Sequence Repeats (SSRs)

MISA-web microsatellite identification tool (https://webblast.ipk-gatersleben.de/misa/)^75^ was used to identify SSRs with the following default parameters: the minimum number of repeats for SSR motif of mono-, di-, tri-, tetra-, penta-, hexa-was set to 10, 6, 5, 5, 5, and 5, respectively; the maximum length of sequence between two distinct SSRs was 100 bp.

### Long Repeat Analysis

Web-based REPuter (https://bibiserv.cebitec.uni-bielefeld.de/reputer/)^76^ was used to identify forward, reverse, complement, and palindromic repeat sequences using the default setting of 50 for maximum computed repeats; hamming distance was set to 3, and minimal repeat size was set to 30 bp^76^.

### Codon Usage Analysis

Codon usage and relative synonymous codon usage (RSCU) value of all the annotated protein-coding genes presented in the plastomes of *Garcinia* species were analyzed using the MEGA X software v10.2.1^77^. The RSCU with value >1.00 refers to codon that is frequently used, whereas RSCU with value <1.00 refers to codon that is less frequently used. There is no codon usage bias when the RSCU value = 1.00^78^.

### Plastomes Sequence Alignment and Comparative Analysis

Plastomes alignment and visualization were performed using the online comparison tool mVista in LAGAN mode^79,80^. Mesta was used as a reference for alignment. The inverted repeat (IR) regions and the junction sites of the large single-copy (LSC) and small single-copy (SSC) regions of all the *Garcinia* species were compared using IRscope online webtool^81^ for the visualization of the expansion or contraction events. For both analyses, *Erythroxylum novogranatense*, from Erythroxylaceae family of the same order Malpighiales was included.

### Phylogenomic Analysis

For phylogenomic analysis, sixteen species were used: six *Garcinia* species (including *G. mangostana* var. Manggis, Mesta, and Thailand), five *Populus* species, two *Viola* species, *Erythroxylum novogranatense*, and *Jatropha curcas* from the order Malpighiales and *Arabidopsis thaliana* from the order Brassicales. A total of 74 protein-coding genes (Table S9) that were commonly found in the plastomes of these 16 species (three varieties from *G. mangsotana*) were downloaded from NCBI Organelle Genome database. These protein-coding genes were concatenated before aligned using MAFFT version 7 online tool (https://mafft.cbrc.jp/alignment/server/)^82^. Next, ModelTest-NG v0.1.6^83^ was used to select the DNA Evolutionary Models. The best model selected was GTR+I+G4 and it was used in the subsequent maximum likelihood (ML) analysis using RAxML-NG v1.0.2 tool^84^.

## Data Availability Statement

The complete plastome sequences and *ITS* gene sequences of *Garcinia mangostana* var. Mesta and var. Manggis have been submitted to GenBank. The accession numbers for plastome sequences are MZ823408 and OK572535 for *Garcinia mangostana* var. Mesta and var. Manggis, respectively. Meanwhile, the *ITS* gene sequences of *Garcinia mangostana* var. Mesta and var. Manggis are OK576276 and OK576274, respectively.

## Acknowledgments

We would like to acknowledge the support of this research by Universiti Kebangsaan Malaysia (UKM) Research University grant AP-2012-018. The group is currently supported by UKM Research University grant DIP-2020-005 (H-HG) and NIG-JOINT grant 2021 (2A2021) (NY and H-HG), Japan.

## Author Contribution Statement

C.C.W. and H.H.G. conceived and planned the experiment. C.C.W. wrote the paper and analyzed the data. Y.N. contributed to the data analysis. H.H.G., N.A.N.M., V.K.S., M.A., and Y.N. reviewed and edited the manuscript. All authors read and approved the manuscript.

## Additional information

Supplementary Tables: Figure S1-S13

Supplementary Figures: Table S1-S13

Supplementary Data

