## Supplementary material for "Plastomes of *Garcinia mangostana* L. and comparative analysis with other *Garcinia* species": Supplementary Data.docx

**Supplementary Data 1**. Adjusted ORF of plastome genes from different *Garcinia* species.

>G. anomala_rps19

GTGTCACGTTCACTAAAAAAAAATCCTTTTGTAGCAATTCATTTATTAAGAAAAATAAAT

AAGCTTAACACAAAAGAAGAAAAAGAAATAATAGTAACATGGTCACGAGCATCAACCATT

ATACCTACAATGATTGGCCATACTATTGCTATCCATAATGGAAAAGAACATTTACCTATT

TATATAACAGATCGTATGGTGGGTCATAAATTGGGCGAATTTTCCCCTACTCTAAATTTC

CGGGGACATGCAAAAAATGATAATAAATCTCGTCGATAA

>G. anomala_cemA

ATGAGGCGGGTTCATTTAAAATACGAAAAAATGAAAAAAAAAAACCTTATTTCCCTTCTA

TATTTTATATCTATAATATTGTTGCCCTGGTGGATCTCTTTTTCGTTTAATAAAAGTGTG

GAATCTTGGGTTATTAATTGGTGGAATACTAGTCAATCTGAAATTTTTTTAAATGCTATT

CAAGAAAAGAGTAGTTTAGAAAAATTCATAGAATTAGAGGAACTCTTACTCTTGGACGAA

ATGATAAAGGAATGCCCGGAAACACATCTGCAAAAGTTTATTATCGTAATCCACAAAGAA

ACGATCAAATTGCTCAAAATGTACAATGAGGAGCGTATCTATACTATTTTGCACTTCTCA

ACCAATATAATCTGTTTCGTTATTCTAAGCGGTTATTCGATTCTAGGTAATGAAGAACTT

TTTCTTCTTAATTCTTGGGTTCAAGAATTTCTATATAACCTAAGTGATACAATAAAAGCT

TTTTTTATTCTTTTATTAACCGATTTATGTATAGGATTCCATTCACCCCACGGCTGGGAA

CTACTGATTGGTTCTGTCTACAAGGATTTTGGATTTACTTATAACGATCAAATCATATCT

GGACTTGTTTCCACTTTTCCAGTAATTATCGATACAATTTTAAAATATTGGATCTTCCAT

TATTTAAATCGTGTATCTCCGTCACTTGTAGTGATTTATCATTCAATGAATGACTGA

>G. anomala_ndhA

ATGAATATAACAGACATAATTGATATAATAGAAATTAATTCTTTTTCTAGATTGAGATTG

GAATCTTTAAACGAGATCTATGGAATTCTATGGATGCTTTTCCCTATTTTTATTCTGATA

TTGGGAATCATGATAGGTGTATTAGTAATTGTGTGGTTAGAAAGAGAAATATCTGCAGGG

GTGCAACAACGTATTGGACCTGAGTATGCCGGTCCTTTTGGAGTTCTTCAAGCGCTAGCG

GACGGGACAAAATTACTTTTCAAAGAGAATCTTTTTCCGTCTCGAGGAGATACTCGTTTA

TTCAGTCTTGGACCAGCCATAGCAGTCATATCAACTCTATTAAGCTATTCCGTAATTCCT

TTTAGCTATCACCTTGTTTTAGCTGATCTAACTATTGGTGTTTTTTTATGGATTGCCATT

TCAAGTATTGCCCCTATTGGCCTTCTTATGTCAGGGTATGGATCAAACAATAAATATTCC

TTTTTAGGTGGTCTACGAGCTGCTGCTCAATCAATTAGTTATGAAATACCATTAACTCTC

TGCGTATTATCCATATCTCTATTATCTAACAGTTCAAGTACAGTTGATATAGTTGAAGCA

CAATCAAAATATGGTTTTTGGGGGTGGAATTTTTGGCGTCAACCTATAGGGTTTATCATT

TTTTTAATTTCTTCCTTAGCGGAATGTGAGAGATTGCCCTTTGATTTACCAGAAGCAGAA

GAAGAATTAGTAGCAGGTTATCAAACCGAATATTCGGGTATTAAATTTGGTTTATTTTAT

GTTGCTTCCTATCTAAACTTATTAGTTTCGTCATTATTTGTAACAGTTCTTTACTTGGGT

GGTTGGAATATCTCTATTCCATTCCTTCCTGAATTTTTTGAGCTAACTAAAGTAAATGGA

GTCTGTGGAACAATAATGGGGATCTTTATTACATTAGTTAAAAGTTATTTGTTCTTGTTC

ATTTCTATCACAATAAGATGGACTTTACCTAGACTAAGAATGGATCAACTATTAAATCTT

GGATGGAAATTTCTTTTACCTATATCTCTTGGTAATTTAGTATTAACAACTTCTTTCCAA

TTACTTTCACTCTAA

>G. anomala_ndhD

ACGAATTCTTTTCCTTGGTTAACATTATTTGTAGTTTTACCGATATCCGCGGGTTTTTTA

ATTTTCTTTTTTCCTCATAGAGGTAATAAGGTAATTCGCTGGTATACTATTAGTATTTGT

GTTTTAGAGCTCCTTTTAATGACTTATATATTTTTGTATTATTTTAAATTGGATGATCCA

TTAATACAATTAACAGAGAGTTCTAAATGGATCAATTTTTTGAATTTTTACTGGAGATTG

GGAATAGATGGGATCTCTTTAGGACCCATTTTTTTGACCGGATTTATCACTACTTTAGCT

ACTTTAGCGGCTCGGCCAGTTACCCGGGATTCTCGCTTATTCTATTTTCTGATGTTAGCA

ATGTATAGTGGTCAAATAGGATTATTTTCTTCTCAAGATCTTTTACTTTTTTTCATCATG

TGGGAATTAGAATTAATTCCCGTTTATCTACTTTTATCCATGTGGGGGGGAAAGAAACGT CTATATTCAGCTACAAAGTTTATTTTGTATACTGCAGGAAGCTCCGTTTTTTTATTAATG

GGAGCCTTGGGTATTGCTTTATATGGTTCCAATGAACCAACATTCAATTTTGAAACATCA

GCCAATCAACCATATCCTGCGGTCCTAGAAATATTGTTCTATATTGGATTTCTTATTGCT

TTTGCTGTCAAATCGCCGATTATACCCTTACATACATGGTTACCGGACACCCACGGAGAA

GCACATTACAGTACTTGTATGCTTTTAGCCGGAATCTTATTAAAAATGGGAGCGTATGGA

TTAGTTCGAATCAATATGGAATTGTTACCCCACGCCCATTCTCTATTTTCCCCTTGGTTA

ATAATAGTAGGTGCAATGCAAATAATCTATGCAGCTTCAACATCTTCTGGTCAGCGAAAT

TTAAAAAAAAGAATAGCCTATTCTTCTGTATCTCATATGGGTTTCATAATTATAGGAATT

TACTCTATAAGTGATATGGGACTCAATGGGTCCATTTTACAAATAATATCACATGGATTT

ATTGGCGCTGCACTTTTTTTCTTGGCCGGAACAAGTTATGATAGAATACGTCTTGTTTAT

CTTGACGAAATGGGTGGAATGGCTACTCCAATGCCAAAAATATTCACACTATTCAATATC

TTATCACTAGCTTCCCTTGCATTACCGGGCATGAGTGGTTTTGTTTCGGAATTGATAGTC

TTTTTTGGGATACTTACCACAGAAAAATATCTTTTAATGCCAAAAATAATAATTTCTTTT

GTAATGGCAGTTGGAATGCTATTAACTCCTCTTTATTTATTATCAATGTTACGTCAGATG

TTCTATGGATACAAGTTATTTAATGCCCTAAACTCTTATTGTTTTGATTCTGGGCCGCGG

GAATTATTTGTTTCCATTTCGATCCTTCTGCCTGTAATAAGTATTGGTATTTACCCGGAT

TTTATTTTCTCACTATCAGTTGAGAAAGTCGAAGCTATCATGTCGACCTACTTTTCTAGG

TAA

>G. gummi-gutta_petD

ATGGGAGTAACAAAAAAACCTGACTTGAATGATCCTGTTTTAAGAGCTAAATTGGCTAAG

GGCATGGGTCATAATTATTACGGAGAACCCGCATGGCCGAACGATCTTTTATATATTTTC

CCAGTAGTAATTCTCGGAACTATTGCATGTAATGTAGGATTAGCGGTTCTAGAACCATCA

ATGATTGGCGAACCCGCGGATCCATTTGCAACTCCTTTGGAAATATTGCCTGAATGGTAT

TTCTTTCCTGTATTTCAAATACTTCGTACAGTTCCAAATAAGTTATTGGGTGTTCTTTTA

ATGGTTTCAGTACCCGCAGGATTATTAACAGTACCTTTTTTGGAAAATGTTAATAAATTC

CAAAATCCATTTCGTCGTCCGGTCGCGACAACTGTCTTTTTGATTGGTACCGTAGTGGCC

CTTTGGTTTGGTATTGGAGCAACTTTGCCTATTGATAAATCCTTAACTTTAGGTCTTTTT

CAAATTGATTCAATTGTAAAATCAAATAGCACTACGTATGTATCTAGGGAATAG

>G. gummi-gutta_clpP

ATGCCTATTGGTGTCCCAAAAGTTCCTTTTCGAAATCCTGGAGAGGACGATTCCATTTGG

ATTGACGTATACAACCGACTTTATCGAGAAAGATTACTTTTTTTAGGTCAAGATGTTGAT

AGCGAAATCTCAAATCAACTTATTGGGCTTATGGTCTATCTCAGTATAGAGAGTGAGACC

AAAGATTTGTATTTGTTTATAAACTCTCCTGGCGGATGGGTAATACCTGGAATAGCGATT

TATGATACTATGCAATTTGTGCGACCAGATGTACAAACAGTATGCATGGGATTAGCCGCT

TCAATGGGATCTTTTATCCTGGCCGGGGGAAAAATTACCAAACGTCTAGCATTCCCTCAC

GCTAGGGTAATGATCCATCAACCTATTGCTGGTTTTTATGAGGCACAAATAGGAGAATTT

GTCCTGGAAGCGGAAGAGCTACTTAAATTGCGCGAAATCATCACAAGGGTGTATGCTCAA

AGAACGGGCAAACCTTTATGGGTTGTATCCGAAGACATGGAAAGGGATGTTTTTATGTCA

GCAACAGAAGCCCAAGCTCATGGACTTGTTGATCTTGTAGCAGTTACATAA

>G. mangostana var Thailand_cemA

ATGAGGCGGGTTCATTTAAAATTTTCATACGAAAAAATGAAAAAAAAAAAAGCCCTTATT

TCCCTTCTATATTTTACATCTATAATATTGTTGCCCTGGTGGATCTCTTTTTCGTTTAAT

AAAAGTGTGGAATCTTGGGTTATTAATTGGTGGAATACTAGTCAATCTGAAATTTTTTTA

AATGCTATTCAAGAAAAGAGTAGTTTAGAAAAATTCATAGAATTAGAGGAACTCTTACTC

TTGGACGAAATGATAAAGGAATGCCCGGAAACACATCTACAAAAGTTTATTATCGCAATC

CACAAAGAAACGATCAAATTGCTCAAAATGTACAATGAGGAGCGTATCTATACTATTTTG CACTTCTCAACCAATATAATCTGTTTCATTATTCTAAGTGGTTATTCGATTCTAGGTAAT

GAAGAACTTTTTCTTCTTAATTCTTGGGTTCAAGAATTTCTATATAACCTAAGTGATACA

ATAAAAGCTTTTTTTATTCTTTTATTAACCGATTTATGTATAGGATTCCATTCACCCCAC

GGCTGGGAACTACTGATTGGTTCTGTCTACAAGGATTTTGGATTTACTTATAACGATCAA

ATTATATCTGGACTTGTTTCCACTTTTCCAGTAATTATCGATACAATTTTTAAATATTGG

ATCTTCCATTATTTAAATCGTGTATCTCCGTCACTTGTAGTGATTTATCATTCAATGAAT

GACTGA

>G. mangostana var Thailand_petD

ATGGGAGTAACAAAAAAACCTGACTTGAATGATCCTGTTTTAAGAGCTAAATTGGCTAAA

GGCATGGGTCATAATTATTACGGAGAACCTGCATGGCCGAACGATCTTTTATATATTTTC

CCAGTAGTAATTCTGGGGACTATTGCATGTAATGTAGGATTAGCGGTTCTAGAACCATCA

ATGATTGGCGAACCCGCGGATCCATTTGCAACTCCTTTGGAAATATTGCCTGAATGGTAT

TTCTTTCCTGTATTTCAAATACTTCGTACAGTTCCAAATAAGTTATTGGGTGTTCTTTTA

ATGGTTTCAGTACCCGCAGGATTATTAACAGTACCTTTTTTGGAAAATGTTAATAAATTC

CAAAATCCATTTCGTCGTCCGGTCGCGACAACTGTCTTTTTGATTGGTACCGTAGTGGCC

CTTTGGTTAGGTATTGGAGCAACTTTGCCTATTGATAAATCCTTAACTTTAGGTCTTTTT

CAAATTGATTCAATTGTAAAATAA

>G. mangostana var Thailand_clpP

ATGCCTATTGGTGTCCCAAAAGTTCCTTTTCGAAATCCTGGAGAGGACGATTCCATTTGG

ATTGACGTATACAACCGACTTTATCGAGAAAGATTACTTTTTTTAGGTCAAGATGTTGAT

AGCGAAATCTCAAATCAACTTATTGGGCTTATGGTCTATCTCAGTATAGAGAGTGAGACC

AAAGATTTGTATTTGTTTATAAACTCTCCTGGCGGATGGGTAATACCTGGAATAGCTATT

TATGATACTATGCAATTTGTGCGACCAGATGTACAAACAGTATGCATGGGATTAGCCGCT

TCAATGGGATCTTTTATCCTGGCCGGGGGAAAAATTACCAAACGTCTAGCATTCCCTCAC

GCTAGGGTAATGATCCATCAACCTATTGCTGGTTTTTATGAGGCACAAATAGGAGAATTT

GTCCTGGAAGCGGAAGAGCTACTTAAATTGCGCGAAATCATCACAAGGGTGTATGCTCAA

AGAACGGGCAAACCTTTATGGGTTGTATCCGAAGACATGGAAAGGGATGTTTTTATGTCA

GCAACAGAAGCCCAAGCTCATGGACTTGTTGATCTTGTAGCAGTTACATAA

>G. oblongifolia_petD

ATGGGAGTAACAAAAAAACCTGACTTGAATGATCCTGTTTTAAGAGCTAAATTGGCTAAA

GGCATGGGTCATAATTATTACGGAGAACCCGCATGGCCGAACGATCTTTTATATATTTTC

CCAGTAGTAATTCTCGGAACTATTGCATGTAATGTAGGATTAGCGGTTCTAGAACCATCA

ATGATTGGCGAACCCGCGGATCCATTTGCAACTCCTTTGGAAATATTGCCTGAATGGTAT

TTCTTTCCTGTATTTCAAATACTTCGTACAGTTCCAAATAAGTTATTGGGTGTTCTTTTA

ATGGTTTCAGTACCCGCAGGATTATTAACAGTACCTTTTTTGGAAAATGTTAATAAATTC

CAAAATCCATTTCGTCGTCCGGTGGCGACAACTGTCTTTTTGATTGGTACCGTAGCGGCC

CTTTGGTTAGGTATTGGAGCAACTTTGCCTATTGATAAATCCTTAACTTTAGGTCTTTTT

CAAATTGATTCAATTGGAAAATCAAATAGCACTAGGTATGTATCTAGGGAATAG

>G. oblongifolia_clpP

ATGCCTATTGGTGTCCCAAAAGTTCCTTTTCGAAATCCTGGAGAGGACGATTCCATTTGG

ATTGACGTATACAACCGACTTTATCGAGAAAGATTACTTTTTTTAGGTCAAGATGTTGAT

AGCGAAATCTCAAATCAACTTATTGGACTTATGGTCTATCTCAGTATAGAGAGTGAGACC

AAAGATTTGTATTTGTTTATAAACTCTCCTGGCGGATGGGTAATACCTGGAATAGCGATT

TATGATACTATGCAATTTGTGCGACCAGATGTACAAACAGTATGCATGGGATTAGCCGCT

TCAATGGGATCTTTTATCCTGGCCGGGGGAAAAATTACCAAACGTCTAGCATTCCCTCAC GCTAGGGTAATGATCCATCAACCTATTGCTGGTTTTTATGAGGCACAAATAGGAGAATTT

GTCCTGGAAGCGGAAGAGCTACTTAAATTGCGCGAAATCATCACAAGGGTGTATGCTCAA

AGAACGGGCAAACCTTTATGGGTTGTATCCGAAGACATGGAAAGGGATGTTTTTATGTCA

GCAACAGAAGCCCAAGCTCATGGACTTGTTGATCTTGTAGCAGTTACATAA

>G. oblongifolia_rps16

ATGATAAAACTTCGTTTGAAGCGATGTGGTAGAAACCAACGAACCATTTATCGAATCGTT

GCAATTTATGTTCGATCCCGAGCGGGGGGGCGAGATCTTCAGAAAGTGGGTTTTTATGAT

CCGATAAAAAAAAATCGATTTCAATATTAA

>G. paucinervis_rps12

ATGCCAACTATTAAACAACTTATTAGAAACACAAGACAGCCAATCAAAAATGTCACAAAA

TCCCCCGCTCTTGTGGGCTGTCCTCAGCGACGAGGAACGTGTACTAGGGTGTATACTATC

ACCCCCAAAAAACCAAATTCTGCCTTACGTAAAGTAGCCAGAGTACGATTAACCTCTGGA

TTTGAAATCACTGCTTATATACCTGGTATTGGCCATAATTTACAAGAACATTCTGTAGTC

TTAGTAAGAGGGGGAAGGGTTAAGGATTTACCCGGTGTGAGATATCACATTGTTCGAGGA

ACCCTAGATGCTGTCGGAGTAAAGGATCGTCAACAAGGGCGTTCTAAATATGGAGTCAAA

AAGCCAAAATAA

>G. paucinervis_petD

ATGGATTCAACAAAAAAACCTGACTTGAATGATCCTGTTTTAAGAGCTAAATTGGCTAAA

GGCATGGGTCATAATTATTACGGAGAACCTGCATGGCCGAACGATCTTTTATATATTTTC

CCAGTAGTAATTCTCGGAACTATTGCATGTAATGTAGGATTAGCGGTTCTAGAACCATCA

ATGATTGGCGAACCCGCGGATCCATTTGCAACTCCTTTGGAAATATTGCCTGAATGGTAT

TTCTTTCCTGTATTTCAAATACTTCGTACAGTTCCAAATAAGTTATTGGGTGTTCTTTTA

ATGGTTTCAGTACCCGCAGGATTATTAACAGTACCTTTTTTGGAAAATGTTAATAAATTC

CAAAATCCATTTCGTCGTCCGGTCGCGACAACTGTCTTTTTGATTGGTACCGTGGTAGCC

CTTTGGTTAGGTATTGGAGCAACTTTGCCTATTGATAAATCCTTAACTTTGGGTCTTTTT

CAAATTGATTCAATTGTAAAATCAAATAGCACTACGTATGTATCTAGGGAATAG

>G. paucinervis_clpP

ATGCCTATTGGTGTCCCAAAAGTTCCTTTTCGAAATCCGGGAGAGGACGATTCCATTTGG

ATTGACGTATACAACCGACTTTATCGAGAAAGATTACTTTTTTTAGGTCAAGATGTTGAT

AGCGAAATCTCAAATCAACTTATTGGACTTATGGTCTATCTCAGTATAGAGAGTGAGACC

AAAGATTTGTATTTGTTTATAAACTCTCCTGGCGGATGGGTAATACCTGGAATAGCTATT

TATGATACTATGCAATTTGTGCGACCAGATGTACAAACAGTATGCATGGGATTAGCCGCT

TCAATGGGATCTTTTATCCTGGCCGGGGGAAAAATTACCAAACGTCTAGCATTCCCTCAC

GCTAGGGTAATGATCCATCAACCTATTGCTGGTTTTTATGAGGCACAAATAGGAGAATTT

GTCCTGGAAGCGGAAGAGCTACTTAAATTGCGCGAAATCATCACAAGGGTGTATGCTCAA

AGAACGGGCAAACCTTTATGGGTTGTATCCGAAGACATGGAAAGGGATGTTTTTATGTCA

GCAACAGAAGCCCAAGCTCATGGACTTGTTGATCTTGTAGCAGTTACATAA

>G. pedunculata_psbM

ATGGAAGTAAATATTCTCGCCTTTATTGCTACTGCACTCTTCATTCTAGTTCCTACTGCT

TTTTTGCTTATAATATACGTAAAAACTGTTAGTCAAAGCGATTAA

>G. pedunculata_ndhK

ATGAATTCCATTGAGTTTCCCCTACTTGATCGGACAACCCAAACTTCAGTTATTTCAACT ACATCAAATGATCTTTCAAATTGGTCACGACTCTCCAGTTTATGGCCGCTTCTCTATGGT

ACCAGTTGTTGCTTCATTGAATTTGCTGCATTAATAGGCTCACGATTCGACTTTGATCGT

TATGGACTAGTACCAAGATCTAGTCCTAGACAGGCCGACCTTATTTTAACAGCTGGCACA

GTAACCATGAAAATGGCTCCTTCTTTAGTGAGATTATATGAACAAATGCCTGAACCAAAA

TATGTTATTGCTATGGGAGCCTGTACAATTACAGGAGGAATGTTCAGTACCGATTCTTAT

AGTACTGTTCGGGGAGTGGATAAGTTAATTCCTGTCGATGTCTATTTGCCAGGTTGTCCA

CCTAAACCGGAGGCCGTTATAGATGCTATAACAAAACTTCGTAAAAAACTATCTCGAGAA

ATTTATGACGATCGAATTCGGTCCCCACAGGGAAATCAGTGTTTTACTACCAATCATAAG

TTTCATATTGGATGCACTACTCATACCGGAAGTTATGATCAAGGATTGCTCTATCAACCG

CCGACTACTTCCAAAATTCCCCCTGAAACATTTTTCAAATACAAAAAGCCAGTCTCGTCC

TACGAATTAATAAATTAG

>G. pedunculata_ndhE

ATGATGCTCGAACATGTACTTGTTTTGAGTGCCTATTTATTTTCTATCGGTATCTATGGA

TTGATCACGAGTCGAAATATGGTTAGGGCCCTTATGTGCCTTGAACTTATTTTGAATGCT

GTTAATATCAATTTTGTAACATTTTCTGATTTTTTTGATAGTCGACAATTAAAAGGAAAT

ATTTTTTCCATTTTTGTTATAGCTATTGCAGCAGCTGAAGCGGCTATCGGGCTGGCTATT

GTTTCGTCAATTTATCGTAACAGAAAATCCATCCGTATCAATCAATCTAATTTGTTGAAT

AAGTAA

>G. pedunculata_cemA

ATGAGGCGGGTTCATTTAAAATTTTCATACGAAAAAATGAAAAAAAAAAAAGCCCTTATT

TCCCTTCTATATTTTACATCTATAATATTGTTGCCCTGGTGGATCTCTTTTTCGTTTAAT

AAAAGTGTGGAATCTTGGGTTATTAATTGGTGGAATACTAGTCAATCTGAAATTTTTTTT

AATGCTATTCAAGAAAAGAGTAGTTTAGAAAAATTCATAGAATTAGAGGAACTCTTACTC

TTGGACGAAATGATAAAGGAATGCCCGGAAACACATCTACAAAAGTTTATTATCGCAATC

CACAAAGAAACGATCAAATTGCTCAAAATGTACAATGAGGAGCGTATCTATACTATTTTG

CACTTCTCAACCAATATAATCTGTTTCATTATTCTAAGCGGTTATTCGATTCTAGGTAAT

GAAGAACTTTTTCTTCTTAATTCTTGGGTTCAAGAATTTCTATATAACCTAAGTGATACA

ATAAAAGCTTTTTTTATTCTTTTATTAACCGATTTATGTATAGGATTCCATTCACCCCAC

GGCTGGGAACTACTGATTGGTTCTGTCTACAAGGATTTTGGATTTACTTATAACGATCAA

ATTATATCTGGACTTGTTTCCACTTTTCCAGTAATTATCGATACAATTTTAAAATATTGG

ATCTTCCATTATTTAAATCGTGTATCTCCGTCACTTGTAGTGATTTATCATTCAATGAAT

GACTGA

>G. pedunculata_ndhA

ATGAATATAACAGACATAATTGATATAATAGAAATTAATTCTTTTTCTAGATTGAGATTG

GAATCTTTAAACGAGATCTATGGAATTCTATGGATGCTTTTCCCTATTTTTATTCTGATA

TTGGGAATCACGATAGGTGTACTAGTAATTGTGTGGTTAGAAAGAGAAATATCTGCAGGG

GTGCAACAACGTATTGGACCTGAGTATGCCGGTCCTTTTGGAGTTCTTCAAGCGCTAGCG

GACGGGACAAAATTACTTTTTAAAGAGAATCTTTTTCCGTCTCGAGGAGATACTCGTTTA

TTCAGTCTTGGACCAGCCATAGCAGTCATATCAACTCTATTAAGCTATTCCGTAATTCCT

TTTAGCTATCACCTTGTTTTAGCTGATCTAACTATTGGTGTTTTTTTATGGATTGCCATT

TCAAGTATTGCCCCTATTGGCCTTCTTATGTCAGGGTATGGATCAAACAATAAATATTCC

TTTTTAGGTGGTCTACGAGCTGCTGCTCAATCAATTAGTTATGAAATACCATTAACTCTC

TGTGTATTATCCATATCTCTATTATCTAACAGTTCAAGTACAGTTGATATAGTTGAAGCA

CAATCAAAATATGGTTTTTGGGGGTGGAATTTTTGGCGTCAACCTATAGGGTTTATCATT

TTTTTAATTTCTTCCTTAGCGGAATGTGAGAGATTGCCCTTTGATTTACCAGAAGCAGAA

GAAGAATTAGTAGCAGGTTATCAAACCGAATATTCGGGTATTAAATTTGGTTTATTTTAT

GTTGCTTCCTATCTAAACTTATTAGTTTCGTCATTATTTGTAACAGTTCTTTACTTGGGT

GGTTGGAATATCTCTATTCCATTGCTTCCTGAATTTTTTGAGCTAACTAAAGTAAATGGA

GTCTGTGGAACAATAATGGGGATCTTTATTACATTAGTTAAAAGTTATTTGTTCTTGTTC

ATTTCTATCACAATAAGATGGACTTTACCTAGACTAAGAATGGATCAACTATTAAATCTT

GGATGGAAATTTCTTTTACCTATATCTCTTGGTAATTTAGTATTAACAACTTCTTTCCAA

TTACTTTCACTCTAA

>G. pedunculata_ndhD

ACGAATTCTTTTCCTTGGTTAACATTATTTGTAGTTTTCCCGATATCTGCGGGTTTTTTA

ATTTTCTTTTTACCTCATAGAGGTAATAAGGTAATTCGCTGGTATGCTATAAGTATTTCT

ATTTTAGAGCTCCTTTTAATGACTTATATATTTTCGTATTATTTCAAATTGGATGATCCA

TTAATACAATTAACAGAGAGTTCTAAATGGATCAATTTTTTGAATTTTTACTGGAGATTG

GGAATAGATGGGATCTCTTTAGGACCTATTTTTTTGACCGGATTTATCACTACTTTAGCT

ACTTTAGCGGCTCGGCCAGTTACCCGGGATTCTCGCTTATTCTATTTTCTGATGTTAGCA

ATGTATAGTGGTCAAATAGGATTATTTTCTTCTCAAGATCTTTTACTTTTTTTCATCATG

TGGGAATTAGAATTAATTCCCGTTTATCTACTTTTATCCATGTGGGGGGGAAAGAAACGT

CTATATTCAGCTACAAAGTTTATTTTGTATACTGCAGGAAGCTCCGTTTTTTTATTAATG

GGAGCCTTGGGTATTGCTTTATATGGTTCCAATGAACCAACATTCAATTTTGAAACATCA

GCCAATCAACCATATCCCGCGGTCCTAGAAATATTATTCTATATTGGATTTCTTGTTGCT

TTTGCTGTCAAATCGCCGATTATACCCTTACATACATGGTTACCGGACACCCACGGAGAA

GCACATTACAGTACTTGTATGCTTTTAGCCGGAATCTTATTAAAAATGGGAGCGTACGGA

TTAGTTCGAATCAATATGGAATTGTTACCCCACGCCCATTCTATATTTTCCCCTTGGTTA

ATAATAGTAGGTGCAATGCAAATAATCTATGCAGCTTCAACATCTTCTGGTCAGCGAAAT

TTAAAAAAAAGAATAGCCTATTCTTCTGTATCTCATATGGGTTTCATAATTATAGGAATT

TACTCTATAAGTGATATGGGACTCAATGGGGCCATTTTACAAATAATATCACATGGATTT

ATTGGCGCTGCACTTTTTTTCTTGGCCGGAACAAGTTATGATAGAATACGTCTTGTTTAT

CTTGACGAAATGGGCGGAATGGCTACTCCAATGCCAAAAATATTCACACTATTCAATATC

TTATCACTAGCTTCCCTTGCATTACCGGGCATGAGTGGTTTTTTTTCGGAATTGATAGTC

TTTTTGGGGATACTTACCACAGAAAAATATCTTTTAATGTCAAAAATAATAATTTCTTTT

GTAATGGCAGTTGGAATGCTATTAACTCCTCTTTATTTATTATCAATGTTACGTCAGATG

TTCTATGGATACAAGTTATTTAATGGCCTAAACTCTTATTGTTTTGATTCTGGGCCGCGG

GAATTATTTGTTTCCATTTCGATCCTTCTGCCTGTAATAAGTATTGGTATTTACCCGGAT

TTTATTTTCTCACTATCAGTTGAGAAAGTCGAAACTATCATGTTGACCTATTTTTCTAGG

TAA
