## Supplementary material for "Plastomes of *Garcinia mangostana* L. and comparative analysis with other *Garcinia* species": Supplementary Figures.docx

**Figure S1.** Dot plot analysis showing tig00037451_pilon as a circular contig.

**Figure S2.** Read depth of (a) Mesta PacBio subreads and (b) Mesta Illumina clean reads mapped against the Mesta plastome (MZ823408).

**Figure S3.** Read depth of Manggis Illumina clean reads mapped against plastomes of (a) Mesta variety, (b) Mesta*, (c) Thailand variety, (d) Manggis variety, and (e) Manggis*.

**Figure S4.** Sequence alignment result between plastomes of *G. mangostana* var. Manggis and var. Mesta showing the position of gaps and a variable site.

**Figure S5.** Multiple sequence alignment of (a) *infA* gene of *G. anomala* and *G. pedunculata*, (b) *rpl32* gene of *G. pedunculata* with other species.

**Figure S6.** Comparison of amino acid frequency of six *Garcinia* species.

**Figure S7.** Phylogenetic tree (maximum likelihood) construction based on 16 species (three varieties from *G. mangostana*) whole plastome sequence.

**Figure S8.** Phylogenetic tree inferred from the *ITS* gene of *Garcinia* species using Neighbor-Joining method with bootstrap replications set to 1000 (MEGA X Version 10.2.1).

**Figure S9.** Comparison between *G. mangostana* L. (mangosteen) and *G. malaccensis* to identify the substitution sites and indels in internal transcribed spacer (*ITS*) sequences.

**Figure S10.** Positions of *ITS* gene with heterozygosity detected at *ITS* gene of Manggis and Mesta varieties as visualized using IGV.

**Figure S11.** Different methods used in Manggis variety plastome assembly.

**Figure S12.** ClustalW alignment and IGV visualization to confirm the indel and SNP detected in Supplementary Table S12.

**Figure S13.** Example of ORF alignment before and after adjustment.

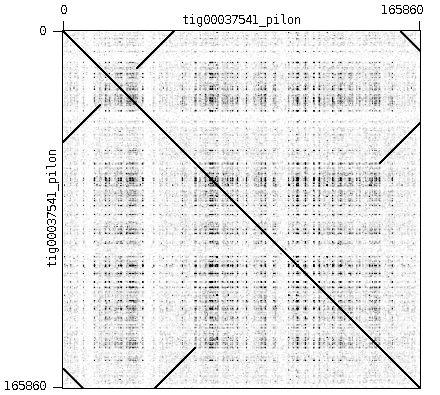

**Figure S1.** Dot plot analysis showing tig00037451_pilon as a circular contig.

**a**

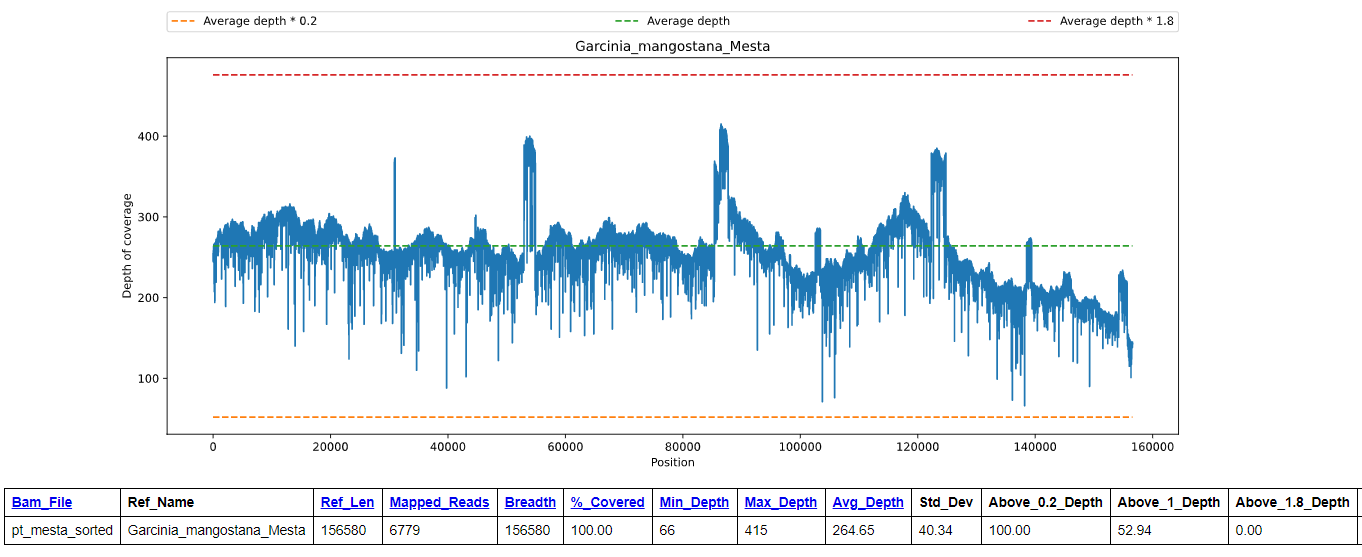

**b**

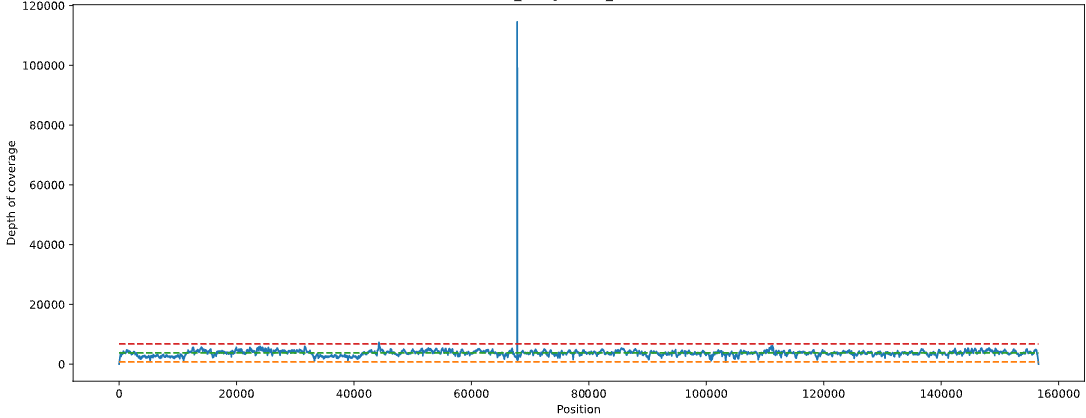

**Figure S2.** Read depth of (a) Mesta PacBio subreads and (b) Mesta Illumina clean reads mapped against the Mesta plastome (MZ823408). * Sharp peak in (b) is due to the TA-rich region.

**a**

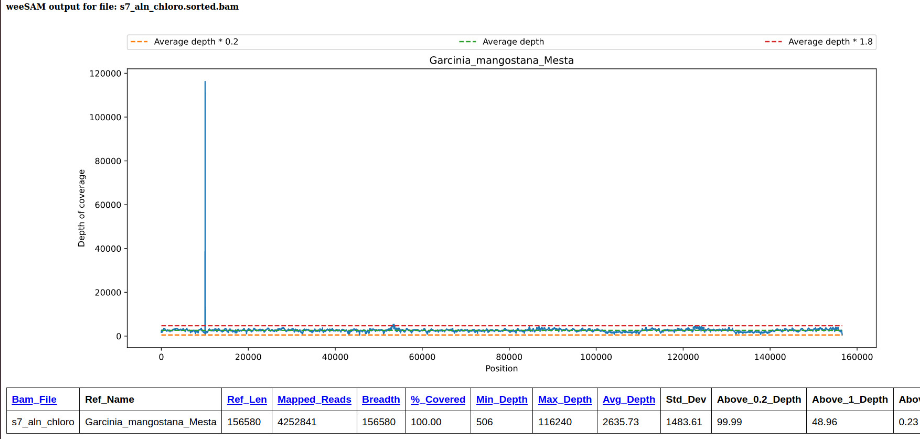

**b**

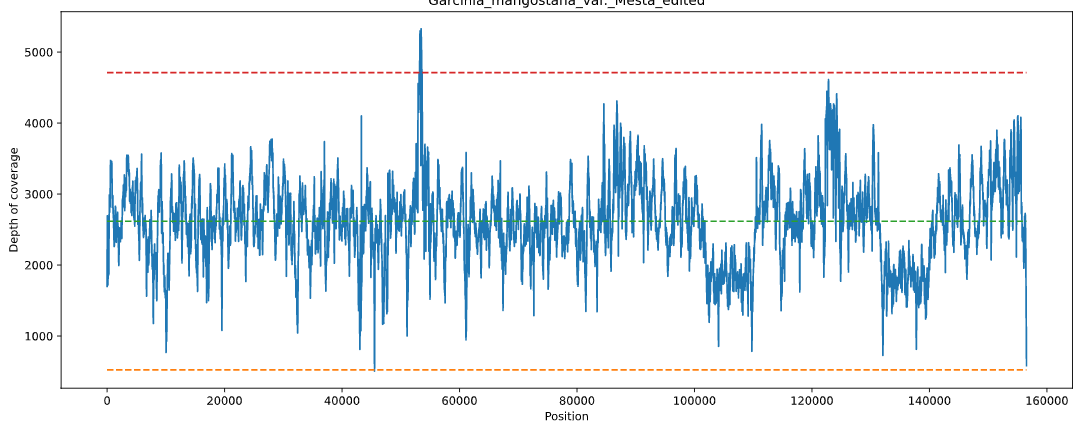

**c**

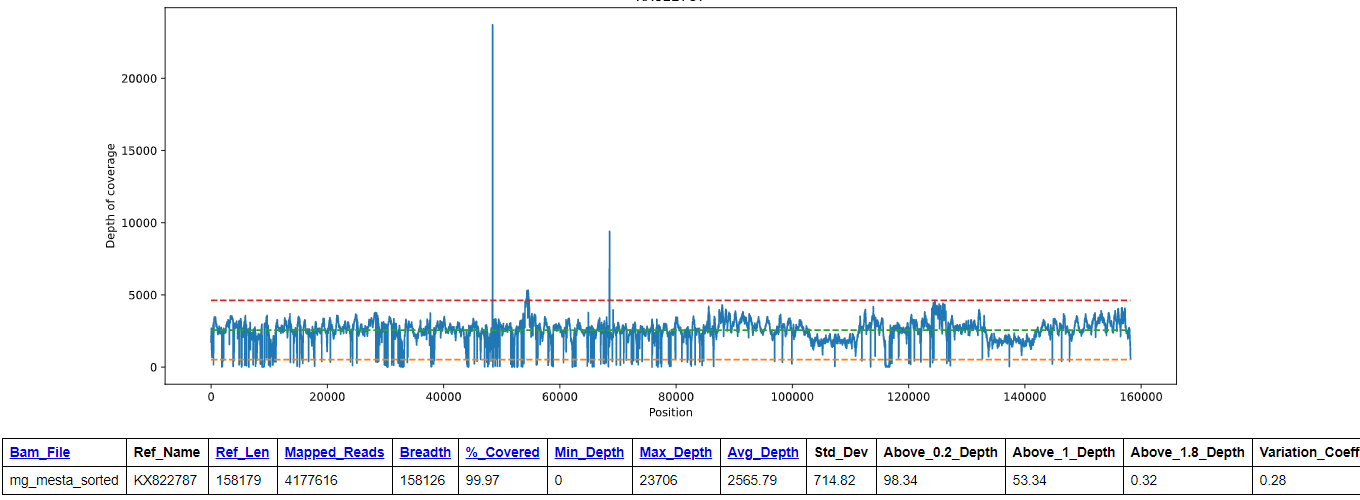

**d**

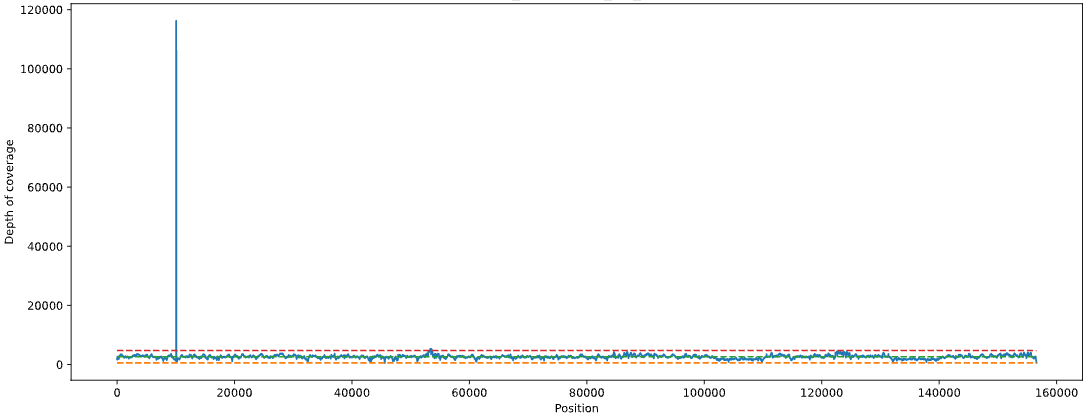

**e**

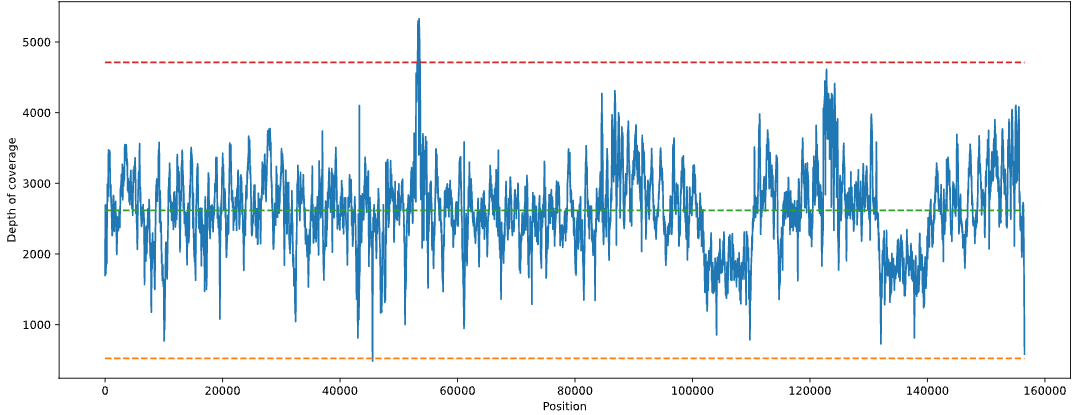

**Figure S3.** Read depth of Manggis Illumina clean reads mapped against plastomes of (a) Mesta variety, (b) Mesta*, (c) Thailand variety, (d) Manggis variety, and (e) Manggis*. *Edited to exclude the TA-rich region of 52 bp.

**Gap**

Position 6,718: Gap1

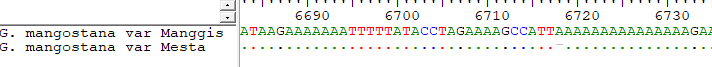

Position 45,577: Gap 2

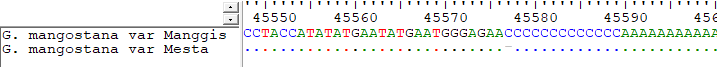

**Variable site**

Position 46,354: Variable site

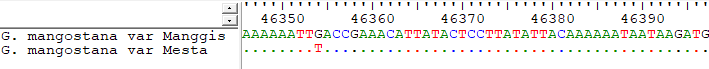

**Figure S4.** Sequence alignment result between plastomes of *G. mangostana* var. Manggis and var. Mesta showing the position of gaps and a variable site.

**a**

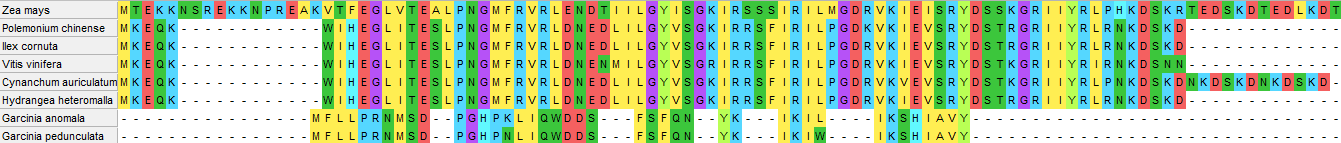

**b**

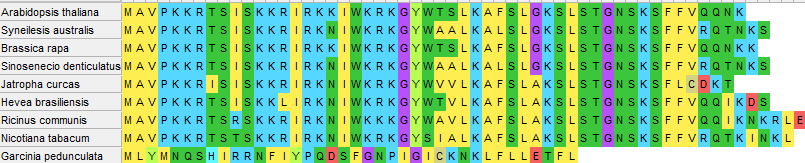

**Figure S5.** Multiple sequence alignment of (a) *infA* gene of *G. anomala* and *G. pedunculata*, (b) *rpl32* gene of *G. pedunculata* with other species.

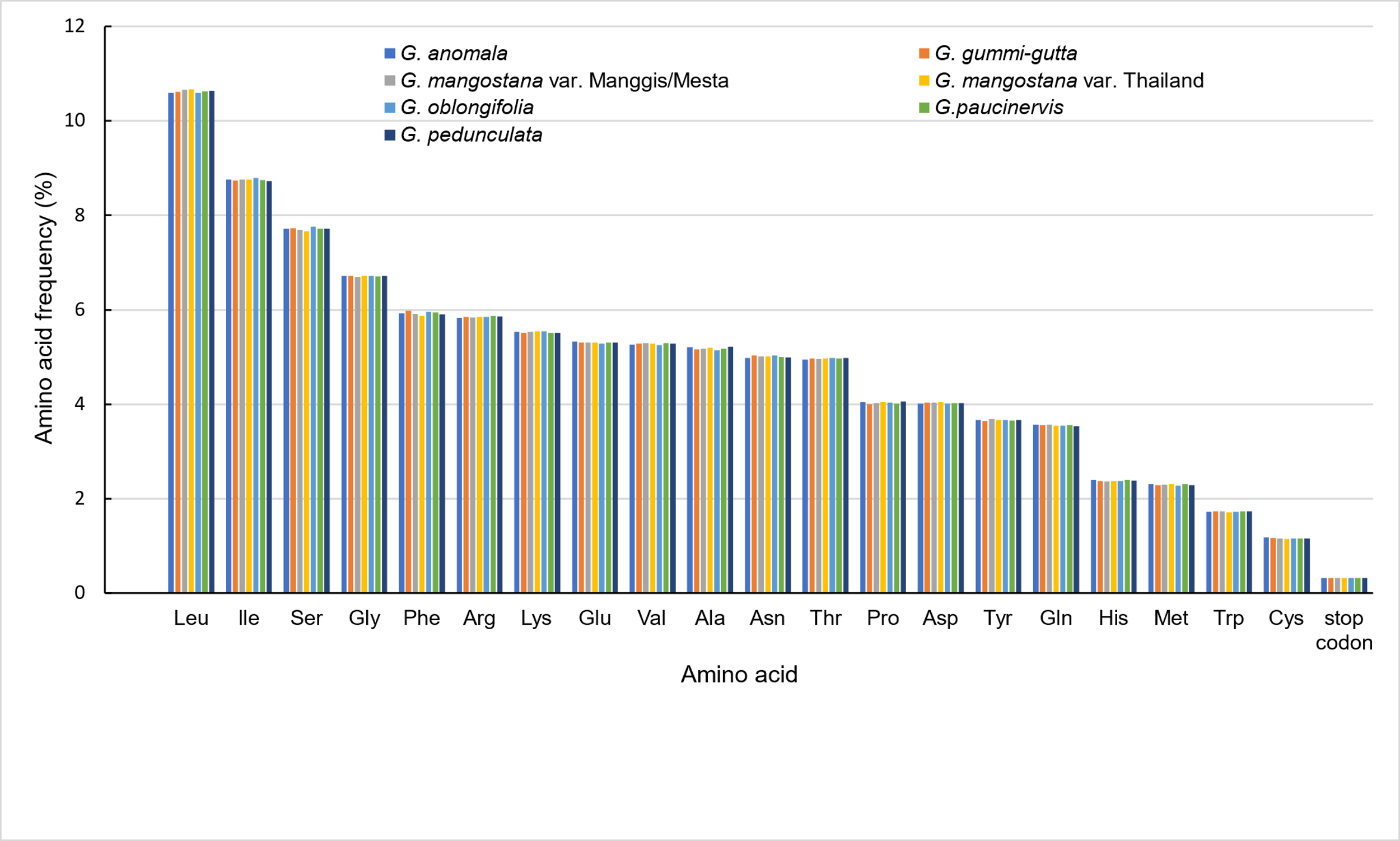

**Figure S6.** Comparison of amino acid frequency of six *Garcinia* species.

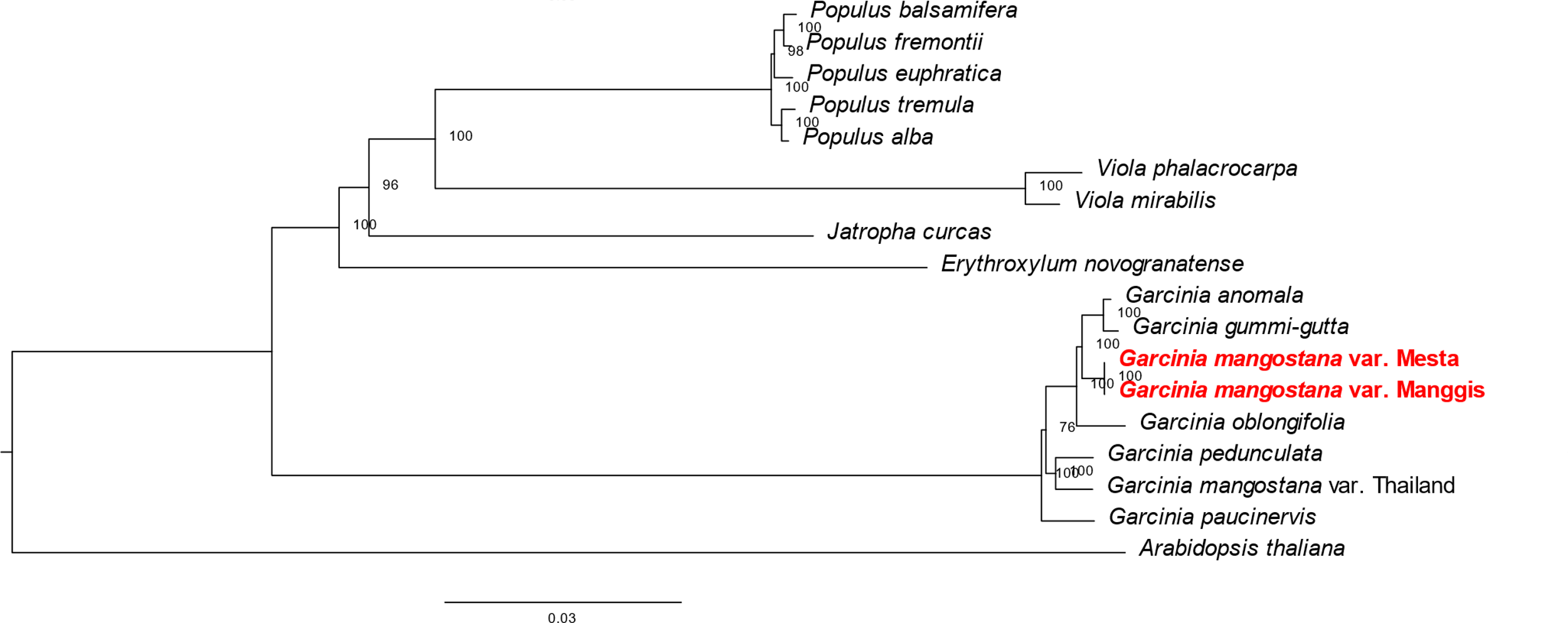

**Figure S7.** Phylogenetic tree (maximum likelihood) construction based on 16 species (three varieties from *G. mangostana*) whole plastome sequence.

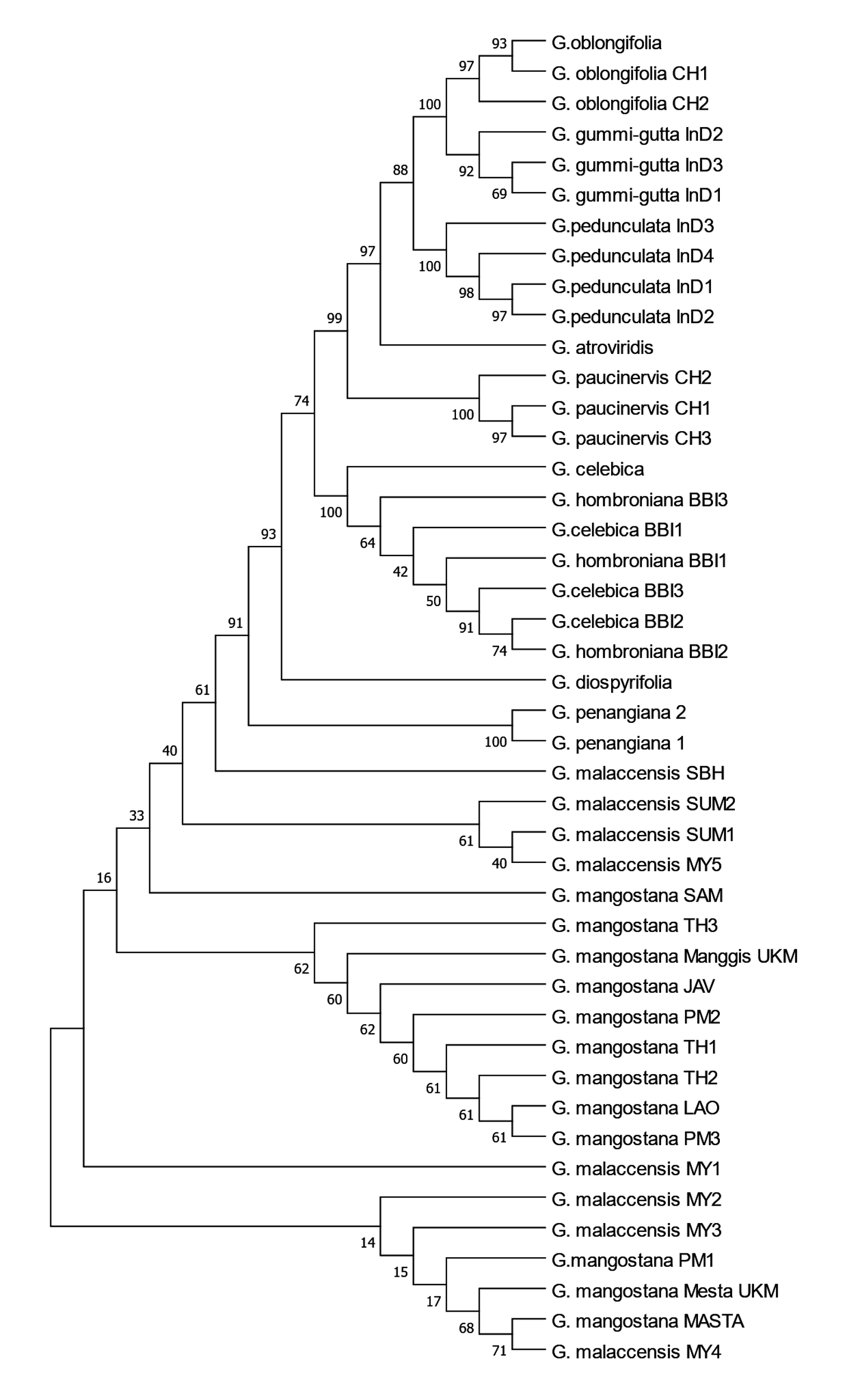

**Figure S8.** Phylogenetic tree inferred from the *ITS* gene of *Garcinia* species using Neighbor-Joining method with bootstrap replications set to 1000 (MEGA X Version 10.2.1). The evolutionary distances were computed using the Kimura 2-parameter method and are in the units of the number of base substitutions per site.

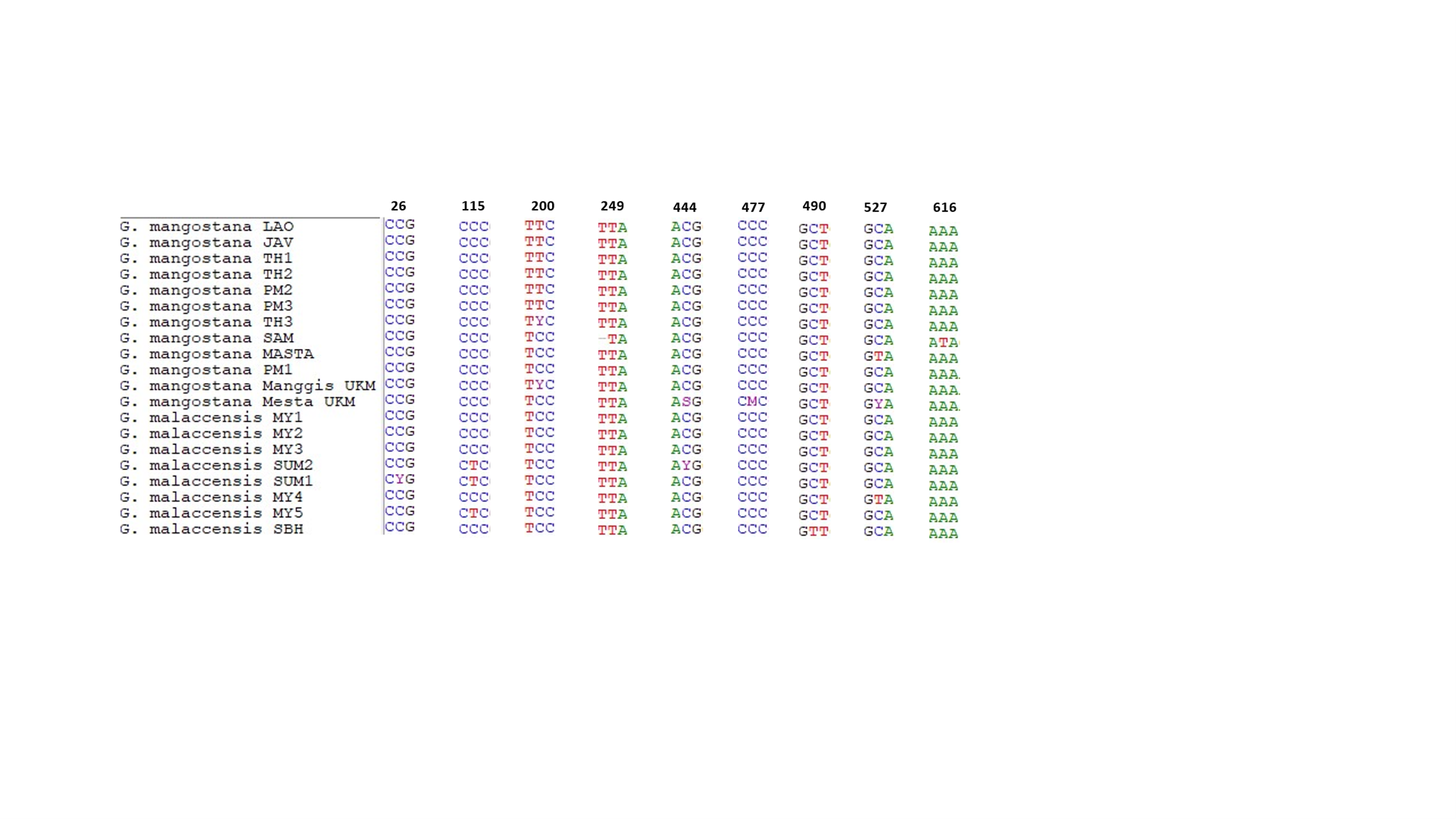

**Figure S9.** Comparison between *G. mangostana* L. (mangosteen) and *G. malaccensis* to identify the substitution sites and indels in internal transcribed spacer (*ITS*) sequences.

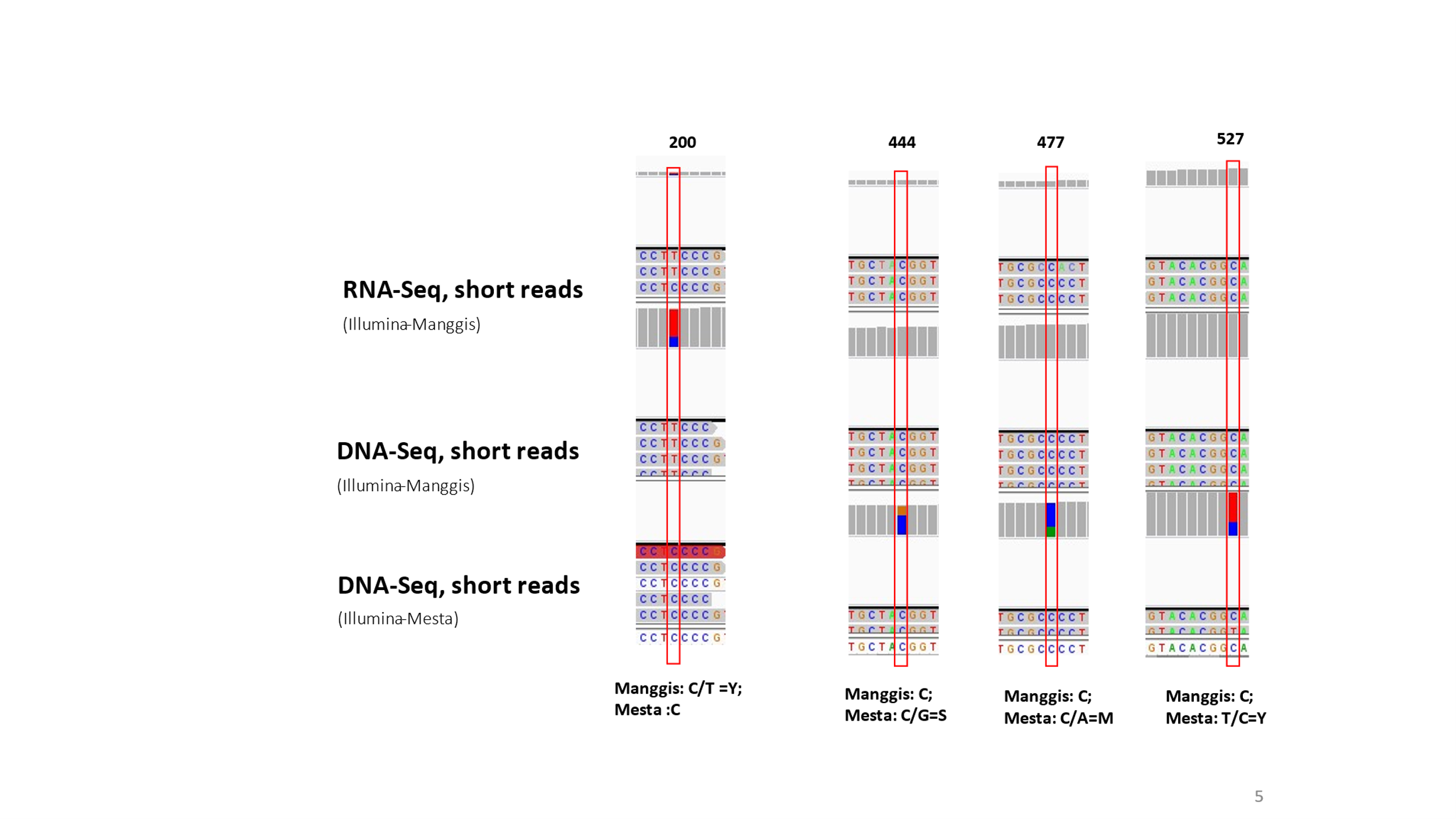

**Figure S10.** Positions of *ITS* gene with heterozygosity detected at *ITS* gene of Manggis and Mesta varieties as visualized using IGV.

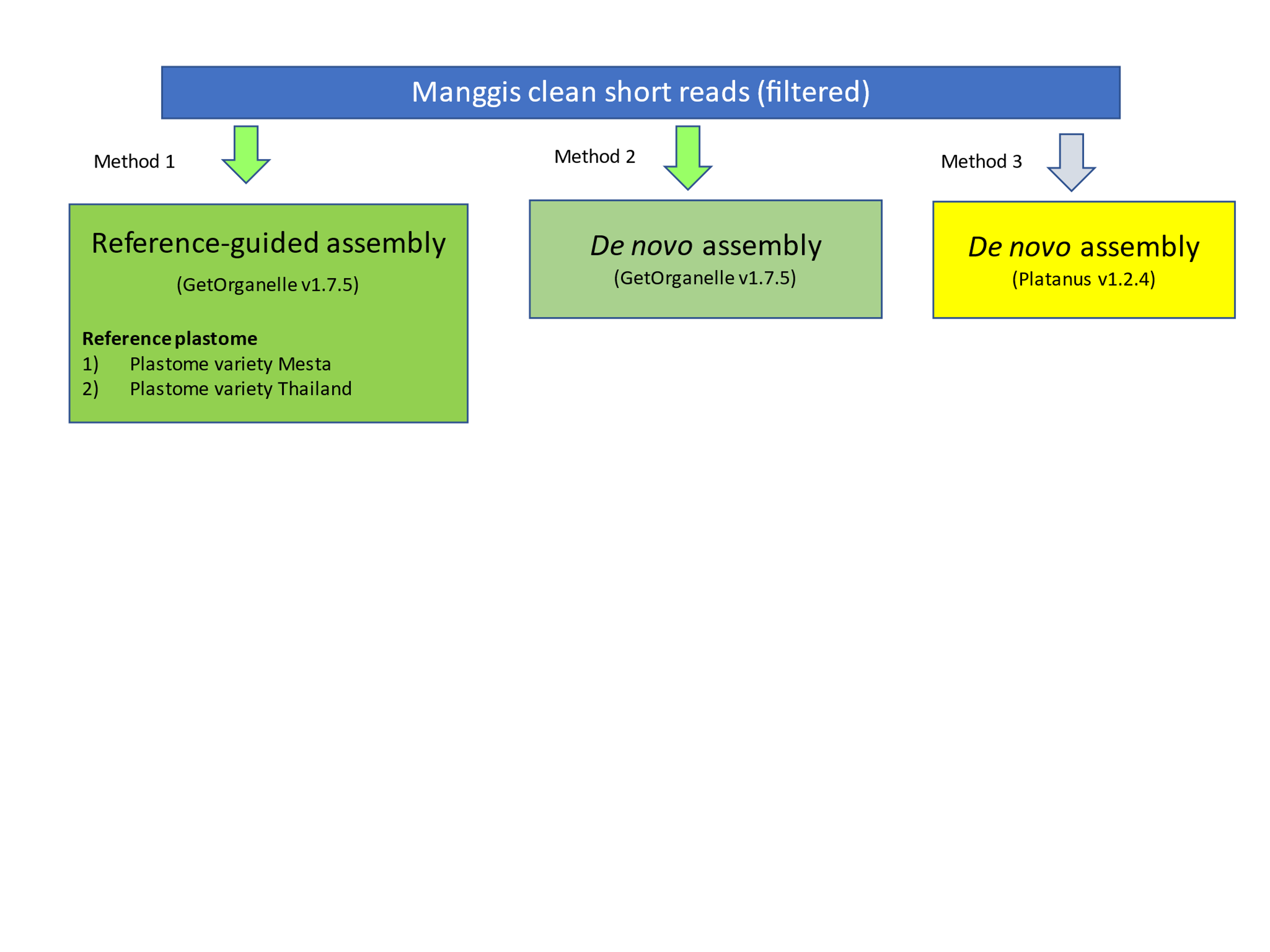

**Figure S11.** Different methods used in Manggis variety plastome assembly.

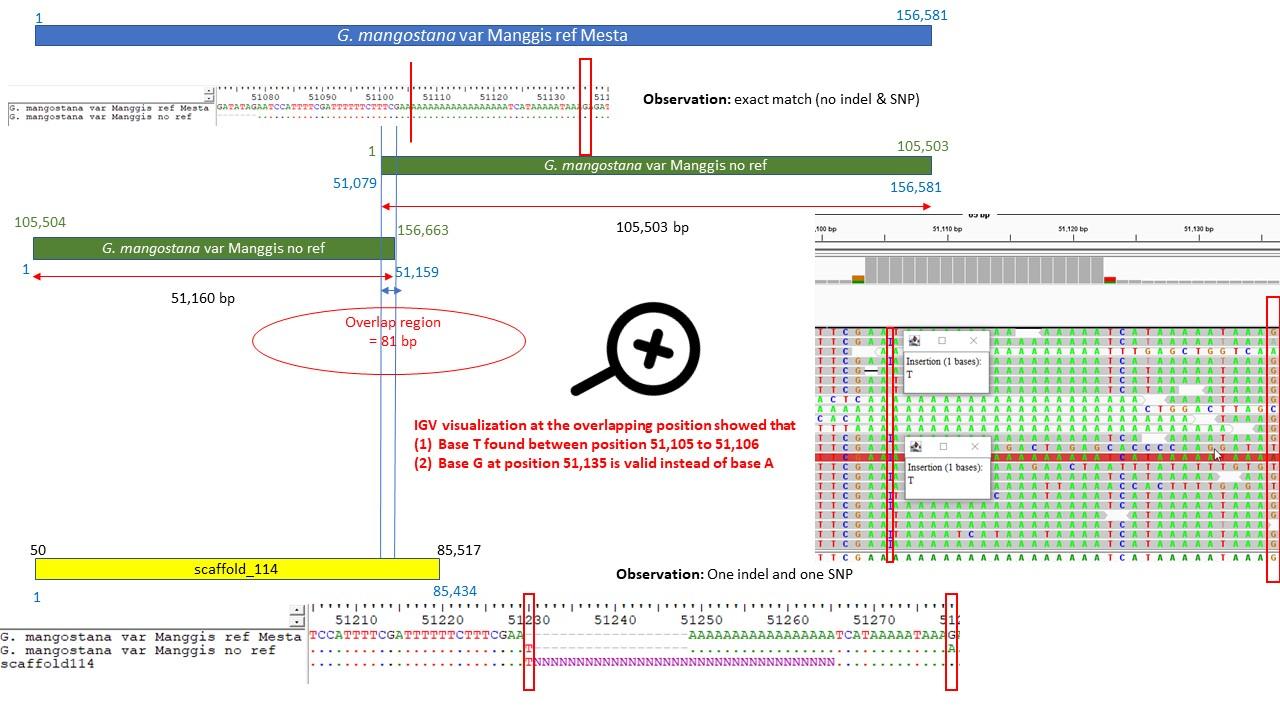

**Figure S12.** ClustalW alignment and IGV visualization to confirm the indel and SNP detected in Supplementary Table S12. ClustalW alignment was performed to detect the position of the one indel and one SNP while IGV tool was used to confirm the existence of indel and SNP. IGV visualization confirmed that base T existed between position of 51,105 and 51,106 while position 51,136 confirmed was base G. Hence, base T was added between position of 51,105 and 51,106 of *G. mangostana* var Manggis ref Mesta.

**Original ORF (*cemA*)**

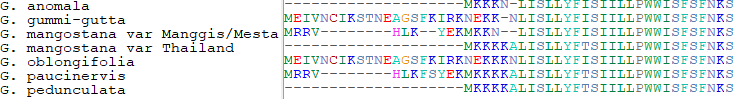

**ORF adjustment at 5’ end (*cemA*)**

**Species with CDS adjusted:** *G. anomala*, *G. mangostana* var Thailand, *G. pedunculata*

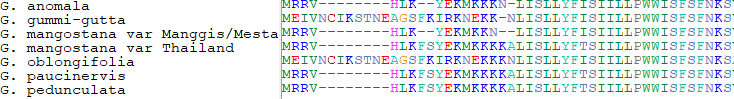

**Original ORF (*petD*)**

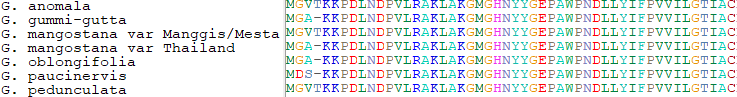

**ORF adjustment at splicing site (*petD*)**

**Species with CDS adjusted:**

*G. gummi-gutta*, *G. mangostana* var Thailand, *G. oblongifolia, G. paucinervis*

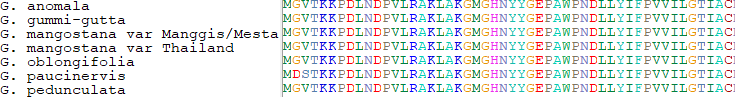

**Original ORF (*rps16*)**

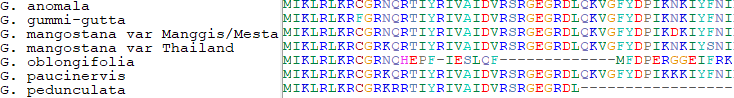

**ORF adjustment at splicing site (*rps16*)**

**Species with CDS adjusted:** *G. oblongifolia*

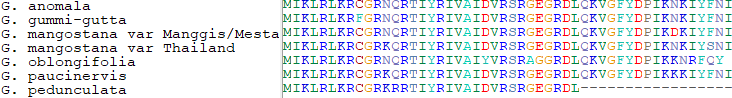

**Figure S13.** Example of ORF alignment before and after adjustment.
