## Supplementary material for "Plastomes of *Garcinia mangostana* L. and comparative analysis with other *Garcinia* species": Supplementary Tables.docx

**Table S1.** Summary statistics of the polished assembled *G. mangostana* var. Mesta genome.

**Table S2.** Summary statistics of the (a) Mesta PacBio subread and (b) Mesta Illumina clean read depth coverage mapped against the Mesta plastome (MZ823408).

**Table S3.** Summary statistics of Manggis Illumina clean read depth coverage against plastomes of Thailand, Mesta, and Manggis varieties.

**Table S4.** Summary of the polymorphism site analysis.

**Table S5.** Genes with intron(s) in the plastomes of *Garcinia* species.

**Table S6.** Relative Synonymous Codon Usage (RSCU) in plastomes of different *Garcinia* species.

**Table S7.** SSRs identified on the plastome of Mesta variety.

**Table S8.** SSRs identified on the plastome of Manggis variety.

**Table S9.** List of protein-coding genes used to construct phylogenomics tree.

**Table S10.** Comparison of polymorphic sites (74 CDS used in phylogenomics tree construction) between *G. mangostana* var Mesta/Manggis versus the other *Garcinia* species.

**Table S11.** List of species used for phylogenetic tree construction using *ITS* gene.

**Table S12.** Summary of different methods used for Manggis plastome assembly.

**Table S13.** CDS length comparison of *Garcinia* species before and after adjustment.

**Table S1.** Summary statistics of the polished assembled *G. mangostana* var. Mesta genome.

| **Features** | ***G. mangostana* var. Mesta** |
| --- | --- |
| **Unfiltered subreads** |  |
| Total number | 1,040,386 |
| Total bases (bp) | 21,364,043,967 |
| **Corrected and Trimmed subreads** |  |
| Total number | 208,217 |
| Total bases (bp) | 914,713,746 |
| **Assembled polished data** |  |
| Total length (bp) | 58,761,678 |
| Number of contigs | 7,616 |
| Largest contig size (bp) | 395,099 |
| N50 (bp) | 10,212 |

**Table S2.** Summary statistics of the (a) Mesta PacBio subread and (b) Mesta Illumina clean-read depth coverage mapped against the Mesta plastome (MZ823408).

| **Sequence data** | **Ref. Length** | **Mapped Reads** | **Breadth** | **% Coverage** | **Min. Depth** | **Max. Depth** | **Avg. Depth** |
| --- | --- | --- | --- | --- | --- | --- | --- |
| PacBio | 156,580 | 6,779 | 156,580 | 100 | 66 | 415 | 265 |
| Illumina | 156,580 | 5,987,076 | 156,580 | 100 | 18 | 114,565 | 3,751 |

**Table S3.** Summary statistics of Manggis Illumina clean data read depth coverage against plastomes of Thailand, Mesta, and Manggis varieties.

| **Variety** | **Length (bp)** | **No. mapped reads** | **Breadth** | **% Coverage** | **Min. Depth** | **Max. Depth** | **Avg. Depth** |
| --- | --- | --- | --- | --- | --- | --- | --- |
| Thailand  KX822787 | 158,179 | 4,177,616 | 158,126 | 99.97 | 0 | 23,706 | 2,566 |
| Mesta | 156,580 | 4,252,841 | 156,580 | 100 | 506 | 116,240 | 2,638 |
| Mesta* | 156,528 | 4,140,697 | 156,528 | 100 | 506 | 5,328 | 2,617 |
| Manggis | 156,582 | 4,252,794 | 156,582 | 100 | 484 | 116,240 | 2,636 |
| Manggis* | 156,530 | 4,140,650 | 156,530 | 100 | 484 | 5,328 | 2,617 |

*Edited to exclude the TA-rich region of 52 bp.

**Table S4.** Summary of the polymorphism site analysis.

| **Features** | **Polymorphism site between plastomes of Manggis and Mesta varieties** |
| --- | --- |
| Number of sites | 156,582 |
| Sites with alignment gaps or missing data | 2 |
| Invariable (monomorphic) sites | 156,579 |
| Variable (polymorphic) sites | 1 |

**Table S5.** Genes with intron(s) in the plastomes of *Garcinia* species. Gene *rps12** is a trans-spliced gene with 5’end located at the LSC regions while the duplicated 3’ ends located at the IR regions.

| **Species** | **Gene** | **Location** | **Exon I (bp)** | **Intron I (bp)** | **Exon II (bp)** | **Intron II (bp)** | **Exon III (bp)** |
| --- | --- | --- | --- | --- | --- | --- | --- |
| *G. anomala* | *rps16* | LSC | 40 | 905 | 179 |  |  |
|  | *atpF* | LSC | 145 | 761 | 398 |  |  |
|  | *rpoC1* | LSC | 432 | 761 | 1632 |  |  |
|  | *ycf3* | LSC | 126 | 717 | 387 |  |  |
|  | *rps12* | LSC | 114 |  | 232 | 538 | 26 |
|  | *clpP* | LSC | 71 | 753 | 292 | 613 | 228 |
|  | *petB* | LSC | 6 | 852 | 642 |  |  |
|  | *petD* | LSC | 8 | 820 | 526 |  |  |
|  | *rpl16* | LSC | 9 | 1172 | 399 |  |  |
|  | *rpl2* | IR | 400 | 661 | 434 |  |  |
|  | *ndhB* | IR | 777 | 699 | 756 |  |  |
|  | *ndhA* | SSC | 561 | 1134 | 534 |  |  |
|  | *trnK-UUU* | LSC | 37 | 2564 | 35 |  |  |
|  | *trnG-UCC* | LSC | 23 | 703 | 48 |  |  |
|  | *trnL-UAA* | LSC | 35 | 644 | 50 |  |  |
|  | *trnV-UAC* | LSC | 39 | 610 | 35 |  |  |
|  | *trnI-GAU* | IR | 37 | 937 | 35 |  |  |
|  | *trnA-UGC* | IR | 38 | 803 | 35 |  |  |
| *G. gummi-gutta* | *rps16* | LSC | 40 | 910 | 179 |  |  |
|  | *atpF* | LSC | 145 | 760 | 398 |  |  |
|  | *rpoC1* | LSC | 432 | 769 | 1632 |  |  |
|  | *ycf3* | LSC | 126 | 718 | 387 |  |  |
|  | *rps12* | LSC | 114 |  | 232 | 538 | 26 |
|  | *clpP* | LSC | 71 | 748 | 292 | 613 | 228 |
|  | *petB* | LSC | 6 | 851 | 642 |  |  |
|  | *petD* | LSC | 8 | 812 | 526 |  |  |
|  | *rpl16* | LSC | 9 | 1152 | 399 |  |  |
|  | *rpl2* | IR | 400 | 661 | 434 |  |  |
|  | *ndhB* | IR | 777 | 699 | 756 |  |  |
|  | *ndhA* | SSC | 562 | 1136 | 533 |  |  |
|  | *trnK-UUU* | LSC | 37 | 2589 | 35 |  |  |
|  | *trnG-UCC* | LSC | 23 | 703 | 48 |  |  |
|  | *trnL-UAA* | LSC | 35 | 619 | 50 |  |  |
|  | *trnV-UAC* | LSC | 39 | 606 | 35 |  |  |
|  | *trnI-GAU* | IR | 37 | 944 | 35 |  |  |
|  | *trnA-UGC* | IR | 38 | 804 | 35 |  |  |

**Table S5 (continue).** Genes with intron(s) in the plastomes of *Garcinia* species. Gene *rps12** is a trans-spliced gene with 5’end located at the LSC regions while the duplicated 3’ ends located at the IR regions.

| **Species** | **Gene** | **Location** | **Exon I (bp)** | **Intron I (bp)** | **Exon II (bp)** | **Intron II (bp)** | **Exon III (bp)** |
| --- | --- | --- | --- | --- | --- | --- | --- |
| *G.*  *mangostana*  var.  Manggis/  Mesta | *rps16* | LSC | 40 | 908 | 179 |  |  |
|  | *atpF* | LSC | 145 | 751 | 398 |  |  |
|  | *rpoC1* | LSC | 432 | 755 | 1635 |  |  |
|  | *ycf3* | LSC | 126 | 722 | 387 |  |  |
|  | *rps12* | LSC | 114 |  | 232 | 538 | 26 |
|  | *clpP* | LSC | 71 | 755 | 292 | 612 | 228 |
|  | *petB* | LSC | 6 | 838 | 642 |  |  |
|  | *petD* | LSC | 8 | 798 | 526 |  |  |
|  | *rpl16* | LSC | 9 | 1217 | 399 |  |  |
|  | *rpl2* | IR | 400 | 661 | 434 |  |  |
|  | *ndhB* | IR | 777 | 697 | 756 |  |  |
|  | *ndhA* | SSC | 562 | 1157 | 533 |  |  |
|  | *trnK-UUU* | LSC | 37 | 2543 | 35 |  |  |
|  | *trnG-UCC* | LSC | 23 | 706 | 48 |  |  |
|  | *trnL-UAA* | LSC | 35 | 624 | 50 |  |  |
|  | *trnV-UAC* | LSC | 39 | 607 | 35 |  |  |
|  | *trnI-GAU* | IR | 37 | 945 | 35 |  |  |
|  | *trnA-UGC* | IR | 38 | 803 | 35 |  |  |
| *G.*  *mangostana*  var.  Thailand | *rps16* | LSC | 40 | 923 | 179 |  |  |
|  | *atpF* | LSC | 145 | 743 | 398 |  |  |
|  | *rpoC1* | LSC | 432 | 753 | 1632 |  |  |
|  | *ycf3* | LSC | 126 | 725 | 387 |  |  |
|  | *rps12* | LSC | 114 |  | 232 | 538 | 26 |
|  | *clpP* | LSC | 71 | 736 | 292 | 637 | 228 |
|  | *petB* | LSC | 6 | 829 | 642 |  |  |
|  | *petD* | LSC | 8 | 813 | 496 |  |  |
|  | *rpl16* | LSC | 9 | 1188 | 399 |  |  |
|  | *rpl2* | IR | 400 | 661 | 434 |  |  |
|  | *ndhB* | IR | 777 | 698 | 756 |  |  |
|  | *ndhA* | SSC | 562 | 1157 | 533 |  |  |
|  | *trnK-UUU* | LSC | 37 | 2560 | 35 |  |  |
|  | *trnG-UCC* | LSC | 23 | 696 | 48 |  |  |
|  | *trnL-UAA* | LSC | 35 | 658 | 50 |  |  |
|  | *trnV-UAC* | LSC | 39 | 596 | 35 |  |  |
|  | *trnI-GAU* | IR | 37 | 944 | 35 |  |  |
|  | *trnA-UGC* | IR | 38 | 795 | 35 |  |  |
|  | *trnI-GAU* | IR | 37 | 941 | 35 |  |  |
|  | *trnA-UGC* | IR | 38 | 804 | 35 |  |  |

**Table S5 (continue).** Genes with intron(s) in the plastomes of *Garcinia* species. Gene *rps12** is a trans-spliced gene with 5’end located at the LSC regions while the duplicated 3’ ends located at the IR regions.

| **Species** | **Gene** | **Location** | **Exon I (bp)** | **Intron I (bp)** | **Exon II (bp)** | **Intron II (bp)** | **Exon III (bp)** |
| --- | --- | --- | --- | --- | --- | --- | --- |
| *G. oblongifolia* | *rps16* | LSC | 40 | 908 | 110 |  |  |
|  | *atpF* | LSC | 145 | 761 | 398 |  |  |
|  | *rpoC1* | LSC | 432 | 758 | 1632 |  |  |
|  | *ycf3* | LSC | 126 | 722 | 387 |  |  |
|  | *rps12* | LSC | 114 |  | 232 | 538 | 26 |
|  | *clpP* | LSC | 71 | 751 | 292 | 629 | 228 |
|  | *petB* | LSC | 6 | 828 | 642 |  |  |
|  | *petD* | LSC | 8 | 798 | 526 |  |  |
|  | *rpl16* | LSC | 9 | 1200 | 399 |  |  |
|  | *rpl2* | IR | 400 | 669 | 434 |  |  |
|  | *ndhB* | IR | 777 | 700 | 756 |  |  |
|  | *ndhA* | SSC | 562 | 1086 | 533 |  |  |
|  | *trnK-UUU* | LSC | 37 | 2585 | 35 |  |  |
|  | *trnG-UCC* | LSC | 23 | 690 | 48 |  |  |
|  | *trnL-UAA* | LSC | 35 | 638 | 50 |  |  |
|  | *trnV-UAC* | LSC | 39 | 601 | 35 |  |  |
|  | *trnI-GAU* | IR | 37 | 944 | 35 |  |  |
|  | *trnA-UGC* | IR | 38 | 801 | 35 |  |  |
| *G. paucinervis* | *rps16* | LSC | 40 | 773 | 224 |  |  |
|  | *atpF* | LSC | 145 | 748 | 398 |  |  |
|  | *rpoC1* | LSC | 432 | 753 | 1632 |  |  |
|  | *ycf3* | LSC | 126 | 729 | 387 |  |  |
|  | *rps12* | LSC | 114 |  | 232 | 538 | 26 |
|  | *clpP* | LSC | 71 | 745 | 292 | 641 | 228 |
|  | *petB* | LSC | 6 | 841 | 642 |  |  |
|  | *petD* | LSC | 8 | 22 | 526 |  |  |
|  | *rpl16* | LSC | 9 | 43 | 399 |  |  |
|  | *rpl2* | IR | 400 | 669 | 434 |  |  |
|  | *ndhB* | IR | 777 | 699 | 756 |  |  |
|  | *ndhA* | SSC | 561 | 1148 | 534 |  |  |
|  | *trnK-UUU* | LSC | 37 | 2552 | 35 |  |  |
|  | *trnG-UCC* | LSC | 23 | 717 | 48 |  |  |
|  | *trnL-UAA* | LSC | 35 | 640 | 50 |  |  |
|  | *trnV-UAC* | LSC | 39 | 597 | 35 |  |  |
|  | *trnI-GAU* | IR | 37 | 943 | 35 |  |  |
|  | *trnA-UGC* | IR | 38 | 801 | 35 |  |  |

**Table S5 (continue).** Genes with intron(s) in the plastomes of *Garcinia* species. Gene *rps12** is a trans-spliced gene with 5’end located at the LSC regions while the duplicated 3’ ends located at the IR regions.

| **Species** | **Gene** | **Location** | **Exon I (bp)** | **Intron I (bp)** | **Exon II (bp)** | **Intron II (bp)** | **Exon III (bp)** |
| --- | --- | --- | --- | --- | --- | --- | --- |
| *G. pedunculata* | *rps16* | LSC | 40 | 924 | 62 |  |  |
|  | *atpF* | LSC | 145 | 744 | 398 |  |  |
|  | *rpoC1* | LSC | 432 | 765 | 1632 |  |  |
|  | *ycf3* | LSC | 126 | 722 | 387 |  |  |
|  | *rps12* | LSC | 114 |  | 232 | 538 | 26 |
|  | *clpP* | LSC | 71 | 756 | 292 | 648 | 228 |
|  | *petB* | LSC | 6 | 804 | 642 |  |  |
|  | *petD* | LSC | 8 | 807 | 526 |  |  |
|  | *rpl16* | LSC | 9 | 1185 | 399 |  |  |
|  | *rpl2* | IR | 400 | 661 | 434 |  |  |
|  | *ndhB* | IR | 777 | 701 | 756 |  |  |
|  | *ndhA* | SSC | 561 | 1140 | 534 |  |  |
|  | *trnK-UUU* | LSC | 37 | 2559 | 35 |  |  |
|  | *trnG-UCC* | LSC | 23 | 694 | 48 |  |  |
|  | *trnL-UAA* | LSC | 35 | 637 | 50 |  |  |
|  | *trnV-UAC* | LSC | 39 | 597 | 35 |  |  |
|  | *trnI-GAU* | IR | 37 | 941 | 35 |  |  |
|  | *trnA-UGC* | IR | 38 | 804 | 35 |  |  |

**Table S6.** Relative Synonymous Codon Usage (RSCU) in plastomes of different *Garcinia* species.

| **Codon** | **Amino acid** | ***G. anomala*** | | ***G. gummi-gutta*** | | | ***G. mangostana*** | | | | | | | | | ***G. oblongifolia*** | | | ***G. paucinervis*** | | | ***G. pedunculata*** | | |
| --- | --- | --- | --- | --- | --- | --- | --- | --- | --- | --- | --- | --- | --- | --- | --- | --- | --- | --- | --- | --- | --- | --- | --- | --- |
|  |  |  |  |  |  |  | **Manggis** | | | **Mesta** | | | **Thailand** | | |  |  |  |  |  |  |  |  |  |
|  |  | **Count** | **RSCU** | | **Count** | **RSCU** | | **Count** | **RSCU** | | **Count** | **RSCU** | | **Count** | **RSCU** | | **Count** | **RSCU** | | **Count** | **RSCU** | | **Count** | **RSCU** |
| GCU(A) | Ala | 632 | 1.85 | | 627 | 1.85 | | 630 | 1.85 | | 630 | 1.85 | | 633 | 1.86 | | 628 | 1.86 | | 629 | 1.85 | | 633 | 1.85 |
| GCC(A) | Ala | 221 | 0.65 | | 221 | 0.65 | | 221 | 0.65 | | 221 | 0.65 | | 220 | 0.65 | | 223 | 0.66 | | 220 | 0.65 | | 223 | 0.65 |
| GCA(A) | Ala | 372 | 1.09 | | 367 | 1.08 | | 373 | 1.1 | | 373 | 1.1 | | 375 | 1.1 | | 365 | 1.08 | | 369 | 1.09 | | 377 | 1.1 |
| GCG(A) | Ala | 143 | 0.42 | | 141 | 0.42 | | 135 | 0.4 | | 135 | 0.4 | | 133 | 0.39 | | 134 | 0.4 | | 139 | 0.41 | | 133 | 0.39 |
| CGU(R) | Arg | 316 | 1.24 | | 320 | 1.25 | | 322 | 1.26 | | 322 | 1.26 | | 318 | 1.24 | | 321 | 1.25 | | 323 | 1.26 | | 313 | 1.22 |
| CGC(R) | Arg | 95 | 0.37 | | 94 | 0.37 | | 94 | 0.37 | | 94 | 0.37 | | 101 | 0.4 | | 97 | 0.38 | | 92 | 0.36 | | 107 | 0.42 |
| CGA(R) | Arg | 378 | 1.48 | | 380 | 1.48 | | 379 | 1.48 | | 379 | 1.48 | | 371 | 1.45 | | 380 | 1.48 | | 380 | 1.48 | | 371 | 1.45 |
| CGG(R) | Arg | 116 | 0.46 | | 113 | 0.44 | | 113 | 0.44 | | 113 | 0.44 | | 114 | 0.45 | | 111 | 0.43 | | 113 | 0.44 | | 118 | 0.46 |
| AGA(R) | Arg | 468 | 1.84 | | 475 | 1.85 | | 470 | 1.84 | | 470 | 1.84 | | 476 | 1.86 | | 473 | 1.85 | | 477 | 1.86 | | 468 | 1.83 |
| AGG(R) | Arg | 156 | 0.61 | | 155 | 0.61 | | 156 | 0.61 | | 156 | 0.61 | | 154 | 0.6 | | 155 | 0.61 | | 155 | 0.6 | | 159 | 0.62 |
| AAU(N) | Asn | 1029 | 1.57 | | 1044 | 1.58 | | 1035 | 1.57 | | 1035 | 1.57 | | 1034 | 1.57 | | 1042 | 1.58 | | 1032 | 1.57 | | 1031 | 1.58 |
| AAC(N) | Asn | 279 | 0.43 | | 279 | 0.42 | | 281 | 0.43 | | 281 | 0.43 | | 281 | 0.43 | | 279 | 0.42 | | 279 | 0.43 | | 277 | 0.42 |
| GAU(D) | Asp | 843 | 1.6 | | 850 | 1.6 | | 849 | 1.6 | | 849 | 1.6 | | 851 | 1.6 | | 843 | 1.6 | | 845 | 1.6 | | 841 | 1.6 |
| GAC(D) | Asp | 209 | 0.4 | | 210 | 0.4 | | 209 | 0.4 | | 209 | 0.4 | | 210 | 0.4 | | 211 | 0.4 | | 211 | 0.4 | | 212 | 0.4 |
| UGU(C) | Cys | 236 | 1.52 | | 236 | 1.55 | | 237 | 1.56 | | 237 | 1.56 | | 232 | 1.54 | | 237 | 1.56 | | 235 | 1.55 | | 234 | 1.55 |
| UGC(C) | Cys | 74 | 0.48 | | 69 | 0.45 | | 67 | 0.44 | | 67 | 0.44 | | 69 | 0.46 | | 67 | 0.44 | | 69 | 0.45 | | 68 | 0.45 |
| CAA(Q) | Gln | 744 | 1.59 | | 742 | 1.59 | | 743 | 1.59 | | 743 | 1.59 | | 734 | 1.58 | | 741 | 1.59 | | 741 | 1.59 | | 734 | 1.59 |
| CAG(Q) | Gln | 193 | 0.41 | | 191 | 0.41 | | 193 | 0.41 | | 193 | 0.41 | | 196 | 0.42 | | 191 | 0.41 | | 192 | 0.41 | | 192 | 0.41 |
| GAA(E) | Glu | 1075 | 1.54 | | 1072 | 1.54 | | 1074 | 1.54 | | 1074 | 1.54 | | 1071 | 1.54 | | 1068 | 1.54 | | 1073 | 1.54 | | 1076 | 1.55 |
| GAG(E) | Glu | 323 | 0.46 | | 322 | 0.46 | | 318 | 0.46 | | 318 | 0.46 | | 320 | 0.46 | | 321 | 0.46 | | 320 | 0.46 | | 315 | 0.45 |
| GGU(G) | Gly | 543 | 1.23 | | 553 | 1.25 | | 551 | 1.25 | | 551 | 1.25 | | 545 | 1.24 | | 548 | 1.24 | | 552 | 1.25 | | 545 | 1.24 |
| GGC(G) | Gly | 207 | 0.47 | | 196 | 0.44 | | 194 | 0.44 | | 194 | 0.44 | | 204 | 0.46 | | 200 | 0.45 | | 199 | 0.45 | | 203 | 0.46 |
| GGA(G) | Gly | 714 | 1.62 | | 709 | 1.61 | | 714 | 1.62 | | 714 | 1.62 | | 715 | 1.62 | | 718 | 1.63 | | 708 | 1.61 | | 717 | 1.63 |
| GGG(G) | Gly | 298 | 0.68 | | 305 | 0.69 | | 299 | 0.68 | | 299 | 0.68 | | 297 | 0.67 | | 298 | 0.68 | | 301 | 0.68 | | 296 | 0.67 |
| CAU(H) | His | 482 | 1.54 | | 476 | 1.53 | | 475 | 1.53 | | 475 | 1.53 | | 476 | 1.53 | | 479 | 1.54 | | 480 | 1.53 | | 480 | 1.54 |
| CAC(H) | His | 145 | 0.46 | | 147 | 0.47 | | 146 | 0.47 | | 146 | 0.47 | | 145 | 0.47 | | 145 | 0.46 | | 148 | 0.47 | | 145 | 0.46 |
| AUU(I) | Ile | 1152 | 1.5 | | 1148 | 1.5 | | 1153 | 1.5 | | 1153 | 1.5 | | 1154 | 1.51 | | 1155 | 1.5 | | 1151 | 1.5 | | 1140 | 1.5 |
| AUC(I) | Ile | 415 | 0.54 | | 419 | 0.55 | | 422 | 0.55 | | 422 | 0.55 | | 416 | 0.54 | | 423 | 0.55 | | 423 | 0.55 | | 420 | 0.55 |
| AUA(I) | Ile | 732 | 0.96 | | 728 | 0.95 | | 724 | 0.94 | | 724 | 0.94 | | 727 | 0.95 | | 733 | 0.95 | | 723 | 0.94 | | 727 | 0.95 |
| UUA(L) | Leu | 925 | 1.99 | | 923 | 1.99 | | 929 | 1.99 | | 929 | 1.99 | | 927 | 1.99 | | 912 | 1.97 | | 924 | 1.99 | | 918 | 1.98 |
| UUG(L) | Leu | 547 | 1.18 | | 543 | 1.17 | | 543 | 1.16 | | 543 | 1.16 | | 551 | 1.18 | | 546 | 1.18 | | 542 | 1.17 | | 547 | 1.18 |
| CUU(L) | Leu | 617 | 1.33 | | 618 | 1.33 | | 617 | 1.32 | | 617 | 1.32 | | 617 | 1.32 | | 619 | 1.33 | | 616 | 1.33 | | 616 | 1.33 |
| CUC(L) | Leu | 167 | 0.36 | | 172 | 0.37 | | 174 | 0.37 | | 174 | 0.37 | | 169 | 0.36 | | 171 | 0.37 | | 174 | 0.37 | | 173 | 0.37 |
| CUA(L) | Leu | 359 | 0.77 | | 361 | 0.78 | | 365 | 0.78 | | 365 | 0.78 | | 362 | 0.78 | | 368 | 0.79 | | 362 | 0.78 | | 363 | 0.78 |
| CUG(L) | Leu | 167 | 0.36 | | 172 | 0.37 | | 170 | 0.36 | | 170 | 0.36 | | 171 | 0.37 | | 167 | 0.36 | | 171 | 0.37 | | 169 | 0.36 |
| AAA(K) | Lys | 1112 | 1.53 | | 1102 | 1.52 | | 1110 | 1.53 | | 1110 | 1.53 | | 1105 | 1.52 | | 1108 | 1.52 | | 1100 | 1.52 | | 1104 | 1.53 |
| AAG(K) | Lys | 339 | 0.47 | | 345 | 0.48 | | 344 | 0.47 | | 344 | 0.47 | | 347 | 0.48 | | 347 | 0.48 | | 345 | 0.48 | | 340 | 0.47 |
| AUG(M) | Met | 606 | 1 | | 600 | 1 | | 603 | 1 | | 603 | 1 | | 606 | 1 | | 598 | 1 | | 604 | 1 | | 599 | 1 |
| UUU(F) | Phe | 1045 | 1.34 | | 1060 | 1.35 | | 1047 | 1.35 | | 1047 | 1.35 | | 1040 | 1.35 | | 1051 | 1.34 | | 1051 | 1.35 | | 1045 | 1.35 |
| UUC(F) | Phe | 511 | 0.66 | | 510 | 0.65 | | 507 | 0.65 | | 507 | 0.65 | | 500 | 0.65 | | 514 | 0.66 | | 510 | 0.65 | | 500 | 0.65 |
| CCU(P) | Pro | 437 | 1.65 | | 436 | 1.66 | | 437 | 1.66 | | 437 | 1.66 | | 443 | 1.67 | | 436 | 1.65 | | 437 | 1.66 | | 437 | 1.65 |
| CCC(P) | Pro | 189 | 0.71 | | 186 | 0.71 | | 185 | 0.7 | | 185 | 0.7 | | 186 | 0.7 | | 189 | 0.71 | | 186 | 0.71 | | 189 | 0.71 |
| CCA(P) | Pro | 293 | 1.1 | | 289 | 1.1 | | 293 | 1.11 | | 293 | 1.11 | | 285 | 1.08 | | 288 | 1.09 | | 290 | 1.1 | | 295 | 1.11 |
| CCG(P) | Pro | 143 | 0.54 | | 141 | 0.54 | | 141 | 0.53 | | 141 | 0.53 | | 145 | 0.55 | | 146 | 0.55 | | 140 | 0.53 | | 140 | 0.53 |
| UCU(S) | Ser | 553 | 1.64 | | 562 | 1.66 | | 556 | 1.65 | | 556 | 1.65 | | 552 | 1.65 | | 555 | 1.63 | | 559 | 1.66 | | 563 | 1.67 |
| UCC(S) | Ser | 329 | 0.97 | | 323 | 0.95 | | 324 | 0.96 | | 324 | 0.96 | | 326 | 0.97 | | 323 | 0.95 | | 324 | 0.96 | | 325 | 0.96 |
| UCA(S) | Ser | 409 | 1.21 | | 406 | 1.2 | | 404 | 1.2 | | 404 | 1.2 | | 403 | 1.2 | | 413 | 1.22 | | 405 | 1.2 | | 402 | 1.19 |
| UCG(S) | Ser | 201 | 0.59 | | 200 | 0.59 | | 202 | 0.6 | | 202 | 0.6 | | 200 | 0.6 | | 208 | 0.61 | | 200 | 0.59 | | 199 | 0.59 |
| AGU(S) | Ser | 388 | 1.15 | | 394 | 1.16 | | 386 | 1.15 | | 386 | 1.15 | | 386 | 1.15 | | 392 | 1.15 | | 393 | 1.17 | | 387 | 1.15 |
| AGC(S) | Ser | 147 | 0.44 | | 145 | 0.43 | | 148 | 0.44 | | 148 | 0.44 | | 141 | 0.42 | | 147 | 0.43 | | 143 | 0.42 | | 145 | 0.43 |
| ACU(T) | Thr | 543 | 1.67 | | 551 | 1.69 | | 549 | 1.69 | | 549 | 1.69 | | 545 | 1.67 | | 549 | 1.68 | | 551 | 1.69 | | 544 | 1.67 |
| ACC(T) | Thr | 209 | 0.64 | | 201 | 0.62 | | 202 | 0.62 | | 202 | 0.62 | | 207 | 0.64 | | 204 | 0.62 | | 200 | 0.61 | | 207 | 0.63 |
| ACA(T) | Thr | 416 | 1.28 | | 418 | 1.28 | | 417 | 1.28 | | 417 | 1.28 | | 413 | 1.27 | | 417 | 1.28 | | 420 | 1.29 | | 418 | 1.28 |
| ACG(T) | Thr | 129 | 0.4 | | 134 | 0.41 | | 133 | 0.41 | | 133 | 0.41 | | 137 | 0.42 | | 137 | 0.42 | | 132 | 0.41 | | 135 | 0.41 |
| UGG(W) | Trp | 451 | 1 | | 454 | 1 | | 454 | 1 | | 454 | 1 | | 449 | 1 | | 453 | 1 | | 454 | 1 | | 453 | 1 |
| UAU(Y) | Tyr | 802 | 1.67 | | 797 | 1.66 | | 807 | 1.67 | | 807 | 1.67 | | 801 | 1.67 | | 802 | 1.67 | | 798 | 1.66 | | 799 | 1.67 |
| UAC(Y) | Tyr | 160 | 0.33 | | 161 | 0.34 | | 160 | 0.33 | | 160 | 0.33 | | 161 | 0.33 | | 160 | 0.33 | | 161 | 0.34 | | 160 | 0.33 |
| GUU(V) | Val | 514 | 1.49 | | 521 | 1.5 | | 516 | 1.48 | | 516 | 1.48 | | 514 | 1.48 | | 516 | 1.5 | | 523 | 1.51 | | 517 | 1.49 |
| GUC(V) | Val | 172 | 0.5 | | 169 | 0.49 | | 170 | 0.49 | | 170 | 0.49 | | 169 | 0.49 | | 164 | 0.48 | | 168 | 0.48 | | 168 | 0.49 |
| GUA(V) | Val | 501 | 1.45 | | 510 | 1.47 | | 511 | 1.47 | | 511 | 1.47 | | 508 | 1.47 | | 510 | 1.48 | | 508 | 1.46 | | 509 | 1.47 |
| GUG(V) | Val | 195 | 0.56 | | 189 | 0.54 | | 193 | 0.56 | | 193 | 0.56 | | 195 | 0.56 | | 189 | 0.55 | | 191 | 0.55 | | 191 | 0.55 |
| UAA(*) | stop codon | 44 | 1.59 | | 44 | 1.59 | | 44 | 1.59 | | 44 | 1.59 | | 44 | 1.59 | | 43 | 1.55 | | 44 | 1.59 | | 43 | 1.55 |
| UAG(*) | stop codon | 23 | 0.83 | | 22 | 0.8 | | 22 | 0.8 | | 22 | 0.8 | | 22 | 0.8 | | 22 | 0.8 | | 22 | 0.8 | | 24 | 0.87 |
| UGA(*) | stop codon | 16 | 0.58 | | 17 | 0.61 | | 17 | 0.61 | | 17 | 0.61 | | 17 | 0.61 | | 18 | 0.65 | | 17 | 0.61 | | 16 | 0.58 |
| **Total number of codon usage** | | 26,249 | | | 26,265 | | | 26,257 | | | 26,257 | | | 26,216 | | | 26,268 | | | 26,244 | | | 26,195 | |

**Table S7.** SSRs identified on the plastome of Mesta variety.

| No. | Type | SSR | Size | Start | End | Region |
| --- | --- | --- | --- | --- | --- | --- |
| 1 | p1 | (A)13 | 13 | 286 | 298 | LSC |
| 2 | p1 | (A)10 | 10 | 2149 | 2158 | LSC |
| 3 | p1 | (A)14 | 14 | 6718 | 6731 | LSC |
| 4 | c | (T)13atttgaaaatgaaaagattttagattggataagtttaaagacggattttttgtctaccttactttactttaaaactttaaatc(T)12 | 108 | 7111 | 7218 | LSC |
| 5 | p1 | (A)17 | 17 | 7860 | 7876 | LSC |
| 6 | p1 | (A)10 | 10 | 8126 | 8135 | LSC |
| 7 | p1 | (A)10 | 10 | 9141 | 9150 | LSC |
| 8 | p1 | (T)12 | 12 | 9282 | 9293 | LSC |
| 9 | p2 | (TA)12 | 24 | 10060 | 10083 | LSC |
| 10 | p1 | (T)10 | 10 | 10404 | 10413 | LSC |
| 11 | p1 | (A)11 | 11 | 12428 | 12438 | LSC |
| 12 | p1 | (A)12 | 12 | 14994 | 15005 | LSC |
| 13 | p2 | (AT)7 | 14 | 15335 | 15348 | LSC |
| 14 | p1 | (T)12 | 12 | 16420 | 16431 | LSC |
| 15 | p1 | (T)14 | 14 | 17324 | 17337 | LSC |
| 16 | p1 | (T)16 | 16 | 19585 | 19600 | LSC (*rpoC2*) |
| 17 | p1 | (A)10 | 10 | 22222 | 22231 | LSC (*rpoC1*) |
| 18 | c | (A)12gagctactccttactcaagttcccaacgaagaccaagcaccaattcattcttctttgttttgtcca(T)11 | 89 | 23673 | 23761 | LSC |
| 19 | p1 | (T)10 | 10 | 27327 | 27336 | LSC (*rpoB*) |
| 20 | c | (T)11ctcattttttgccc(T)10 | 35 | 30537 | 30571 | LSC |
| 21 | c | (A)10ttaacagtctatttagagtttaatgatttagaatatttgaatttctaatgatatatattatatacaattagtatacaattagaaattcaaaaattga(T)13 | 120 | 32391 | 32510 | LSC |
| 22 | p1 | (T)13 | 13 | 36387 | 36399 | LSC |
| 23 | p1 | (G)10 | 10 | 43078 | 43087 | LSC |
| 24 | c | (T)12cgttttctttaatttctttaaaataaaatatatatatttctttataagagataataagagaaaagaacgaacc(TA)9 | 103 | 43248 | 43350 | LSC |
| 25 | c | (C)12(A)12 | 24 | 45576 | 45599 | LSC |
| 26 | p2 | (TA)7 | 14 | 46972 | 46985 | LSC |
| 27 | p1 | (A)16 | 16 | 47165 | 47180 | LSC |
| 28 | p1 | (A)12 | 12 | 47384 | 47395 | LSC |
| 29 | c | (TA)6attaatataattaatattttattttttattttaattttaggaatatgaaaaaaattgtcttgaatcaatcccaagttcaagaatcagaattg(A)14 | 118 | 47650 | 47767 | LSC |
| 30 | p1 | (T)10 | 10 | 50153 | 50162 | LSC |
| 31 | p1 | (T)12 | 12 | 50955 | 50966 | LSC |
| 32 | p1 | (A)17 | 17 | 51105 | 51121 | LSC |
| 33 | p1 | (T)10 | 10 | 51759 | 51768 | LSC |
| 34 | c | (A)10tgttcgatatcaagtttctcggttaattcaataagaaatcgaagttagtactcgattttgttggtaccatacaacgaattgaattcaa(T)11ctattttgcaaatcagttagttgaacttgaaaattcattgattgaaatag(A)15 | 174 | 54853 | 55026 | LSC |
| 35 | p1 | (T)10 | 10 | 55177 | 55186 | LSC |
| 36 | p1 | (A)10 | 10 | 57049 | 57058 | LSC |
| 37 | c | (T)12aactta(T)10 | 28 | 57579 | 57606 | LSC |
| 38 | p1 | (T)10 | 10 | 60008 | 60017 | LSC |
| 39 | p2 | (AT)8 | 16 | 61164 | 61179 | LSC |
| 40 | c | (T)13acaaaaatttggaattctatctagtgttctagtgttgataagaagactatttgattttatctcttctttcg(T)13ctctaaatctaaattggggggtgattatgtcactattctattgtcagatttaactgttatcgaatgtattaatag(T)10 | 182 | 63643 | 63824 | LSC |
| 41 | p1 | (T)11 | 11 | 66702 | 66712 | LSC |
| 42 | p1 | (A)15 | 15 | 67466 | 67480 | LSC |
| 43 | p3 | (TAT)5 | 15 | 69306 | 69320 | LSC |
| 44 | c | (T)10c(A)11 | 22 | 71353 | 71374 | LSC |
| 45 | p1 | (A)10 | 10 | 72038 | 72047 | LSC |
| 46 | p3 | (TAA)5 | 15 | 72182 | 72196 | LSC |
| 47 | p1 | (T)11 | 11 | 74912 | 74922 | LSC |
| 48 | p1 | (A)11 | 11 | 76296 | 76306 | LSC |
| 49 | p2 | (AT)6 | 12 | 80709 | 80720 | LSC |
| 50 | p1 | (T)10 | 10 | 80921 | 80930 | LSC |
| 51 | p1 | (A)11 | 11 | 82170 | 82180 | LSC |
| 52 | p1 | (A)10 | 10 | 83263 | 83272 | LSC |
| 53 | p1 | (A)11 | 11 | 83984 | 83994 | LSC |
| 54 | p1 | (T)10 | 10 | 85316 | 85325 | LSC (*rps19*) |
| 55 | p1 | (T)11 | 11 | 85638 | 85648 | IRA |
| 56 | p1 | (A)13 | 13 | 91398 | 91410 | IRA (*ycf2*) |
| 57 | p1 | (T)10 | 10 | 98588 | 98597 | IRA |
| 58 | p1 | (T)11 | 11 | 104626 | 104636 | IRA |
| 59 | p1 | (A)12 | 12 | 109667 | 109678 | IRA |
| 60 | p1 | (T)13 | 13 | 109823 | 109835 | IRA |
| 61 | p1 | (A)14 | 14 | 114948 | 114961 | SSC |
| 62 | p1 | (T)13 | 13 | 116651 | 116663 | SSC |
| 63 | p1 | (A)12 | 12 | 121020 | 121031 | SSC |
| 64 | c | (T)13cgttcctcttcttcgttcggaaaaaaagggggcttagcctaaattcgaataaataaagcaaaggattctttcgttcctgatagtca(T)10 | 109 | 122071 | 122179 | SSC |
| 65 | p1 | (T)13 | 13 | 126256 | 126268 | SSC (*ycf1*) |
| 66 | p1 | (T)10 | 10 | 126962 | 126971 | SSC (*ycf1*) |
| 67 | p1 | (T)12 | 12 | 127472 | 127483 | SSC (*ycf1*) |
| 68 | p1 | (T)10 | 10 | 128012 | 128021 | SSC (*ycf1*) |
| 69 | p1 | (A)13 | 13 | 132129 | 132141 | IRB |
| 70 | p1 | (T)12 | 12 | 132286 | 132297 | IRB |
| 71 | p1 | (A)11 | 11 | 137328 | 137338 | IRB |
| 72 | p1 | (A)10 | 10 | 143367 | 143376 | IRB |
| 73 | p1 | (T)13 | 13 | 150554 | 150566 | IRB (*ycf2*) |
| 74 | p1 | (A)11 | 11 | 156316 | 156326 | IRB |
| Gene name in parentheses indicates SSR that is found within the CDS of the respective gene. | | | | | | |
| p1 = mononucleotide; p2 = dinucleotide; p3 = trinucleotide; c = compound microsatellite | | | | | | |

**Table S8.** SSRs identified on the plastome of Manggis variety.

| No. | Type | SSR | Size | Start | End | Region |
| --- | --- | --- | --- | --- | --- | --- |
| 1 | p1 | (A)13 | 13 | 286 | 298 | LSC |
| 2 | p1 | (A)10 | 10 | 2149 | 2158 | LSC |
| 3 | p1 | (A)15 | 15 | 6718 | 6732 | LSC |
| 4 | c | (T)13atttgaaaatgaaaagattttagattggataagtttaaagacggattttttgtctaccttactttactttaaaactttaaatc(T)12 | 108 | 7112 | 7219 | LSC |
| 5 | p1 | (A)17 | 17 | 7861 | 7877 | LSC |
| 6 | p1 | (A)10 | 10 | 8127 | 8136 | LSC |
| 7 | p1 | (A)10 | 10 | 9142 | 9151 | LSC |
| 8 | p1 | (T)12 | 12 | 9283 | 9294 | LSC |
| 9 | p2 | (TA)12 | 24 | 10061 | 10084 | LSC |
| 10 | p1 | (T)10 | 10 | 10405 | 10414 | LSC |
| 11 | p1 | (A)11 | 11 | 12429 | 12439 | LSC |
| 12 | p1 | (A)12 | 12 | 14995 | 15006 | LSC |
| 13 | p2 | (AT)7 | 14 | 15336 | 15349 | LSC |
| 14 | p1 | (T)12 | 12 | 16421 | 16432 | LSC |
| 15 | p1 | (T)14 | 14 | 17325 | 17338 | LSC |
| 16 | p1 | (T)16 | 16 | 19586 | 19601 | LSC (*rpoC2*) |
| 17 | p1 | (A)10 | 10 | 22223 | 22232 | LSC (*rpoC1*) |
| 18 | c | (A)12gagctactccttactcaagttcccaacgaagaccaagcaccaattcattcttctttgttttgtcca(T)11 | 89 | 23674 | 23762 | LSC |
| 19 | p1 | (T)10 | 10 | 27328 | 27337 | LSC (*rpoB*) |
| 20 | c | (T)11ctcattttttgccc(T)10 | 35 | 30538 | 30572 | LSC |
| 21 | c | (A)10ttaacagtctatttagagtttaatgatttagaatatttgaatttctaatgatatatattatatacaattagtatacaattagaaattcaaaaattga(T)13 | 120 | 32392 | 32511 | LSC |
| 22 | p1 | (T)13 | 13 | 36388 | 36400 | LSC |
| 23 | p1 | (G)10 | 10 | 43079 | 43088 | LSC |
| 24 | c | (T)12cgttttctttaatttctttaaaataaaatatatatatttctttataagagataataagagaaaagaacgaacc(TA)9 | 103 | 43249 | 43351 | LSC |
| 25 | c | (C)13(A)12 | 25 | 45577 | 45601 | LSC |
| 26 | p2 | (TA)7 | 14 | 46974 | 46987 | LSC |
| 27 | p1 | (A)16 | 16 | 47167 | 47182 | LSC |
| 28 | p1 | (A)12 | 12 | 47386 | 47397 | LSC |
| 29 | c | (TA)6attaatataattaatattttattttttattttaattttaggaatatgaaaaaaattgtcttgaatcaatcccaagttcaagaatcagaattg(A)14 | 118 | 47652 | 47769 | LSC |
| 30 | p1 | (T)10 | 10 | 50155 | 50164 | LSC |
| 31 | p1 | (T)12 | 12 | 50957 | 50968 | LSC |
| 32 | p1 | (A)17 | 17 | 51107 | 51123 | LSC |
| 33 | p1 | (T)10 | 10 | 51761 | 51770 | LSC |
| 34 | c | (A)10tgttcgatatcaagtttctcggttaattcaataagaaatcgaagttagtactcgattttgttggtaccatacaacgaattgaattcaa(T)11ctattttgcaaatcagttagttgaacttgaaaattcattgattgaaatag(A)15 | 174 | 54855 | 55028 | LSC |
| 35 | p1 | (T)10 | 10 | 55179 | 55188 | LSC |
| 36 | p1 | (A)10 | 10 | 57051 | 57060 | LSC |
| 37 | c | (T)12aactta(T)10 | 28 | 57581 | 57608 | LSC |
| 38 | p1 | (T)10 | 10 | 60010 | 60019 | LSC |
| 39 | p2 | (AT)8 | 16 | 61166 | 61181 | LSC |
| 40 | c | (T)13acaaaaatttggaattctatctagtgttctagtgttgataagaagactatttgattttatctcttctttcg(T)13ctctaaatctaaattggggggtgattatgtcactattctattgtcagatttaactgttatcgaatgtattaatag(T)10 | 182 | 63645 | 63826 | LSC |
| 41 | p1 | (T)11 | 11 | 66704 | 66714 | LSC |
| 42 | p1 | (A)15 | 15 | 67468 | 67482 | LSC |
| 43 | p3 | (TAT)5 | 15 | 69308 | 69322 | LSC |
| 44 | c | (T)10c(A)11 | 22 | 71355 | 71376 | LSC |
| 45 | p1 | (A)10 | 10 | 72040 | 72049 | LSC |
| 46 | p3 | (TAA)5 | 15 | 72184 | 72198 | LSC |
| 47 | p1 | (T)11 | 11 | 74914 | 74924 | LSC |
| 48 | p1 | (A)11 | 11 | 76298 | 76308 | LSC |
| 49 | p2 | (AT)6 | 12 | 80711 | 80722 | LSC |
| 50 | p1 | (T)10 | 10 | 80923 | 80932 | LSC |
| 51 | p1 | (A)11 | 11 | 82172 | 82182 | LSC |
| 52 | p1 | (A)10 | 10 | 83265 | 83274 | LSC |
| 53 | p1 | (A)11 | 11 | 83986 | 83996 | LSC |
| 54 | p1 | (T)10 | 10 | 85318 | 85327 | LSC (*rps19*) |
| 55 | p1 | (T)11 | 11 | 85640 | 85650 | IR |
| 56 | p1 | (A)13 | 13 | 91400 | 91412 | IR (*ycf2*) |
| 57 | p1 | (T)10 | 10 | 98590 | 98599 | IR |
| 58 | p1 | (T)11 | 11 | 104628 | 104638 | IR |
| 59 | p1 | (A)12 | 12 | 109669 | 109680 | IR |
| 60 | p1 | (T)13 | 13 | 109825 | 109837 | IR |
| 61 | p1 | (A)14 | 14 | 114950 | 114963 | SSC |
| 62 | p1 | (T)13 | 13 | 116653 | 116665 | SSC |
| 63 | p1 | (A)12 | 12 | 121022 | 121033 | SSC |
| 64 | c | (T)13cgttcctcttcttcgttcggaaaaaaagggggcttagcctaaattcgaataaataaagcaaaggattctttcgttcctgatagtca(T)10 | 109 | 122073 | 122181 | SSC |
| 65 | p1 | (T)13 | 13 | 126258 | 126270 | SSC (*ycf1*) |
| 66 | p1 | (T)10 | 10 | 126964 | 126973 | SSC (*ycf1*) |
| 67 | p1 | (T)12 | 12 | 127474 | 127485 | SSC (*ycf1*) |
| 68 | p1 | (T)10 | 10 | 128014 | 128023 | SSC (*ycf1*) |
| 69 | p1 | (A)13 | 13 | 132131 | 132143 | IR |
| 70 | p1 | (T)12 | 12 | 132288 | 132299 | IR |
| 71 | p1 | (A)11 | 11 | 137330 | 137340 | IR |
| 72 | p1 | (A)10 | 10 | 143369 | 143378 | IR |
| 73 | p1 | (T)13 | 13 | 150556 | 150568 | IR (*ycf2*) |
| 74 | p1 | (A)11 | 11 | 156318 | 156328 | IR |
| Gene name in parentheses indicates SSR that is found within the CDS of the respective gene. | | | | | | |
| p1 = mononucleotide; p2 = dinucleotide; p3 = trinucleotide; c = compound microsatellite | | | | | | |

**Table S9.** List of protein-coding genes used to construct phylogenomics tree.

| **Protein-coding genes used to construct the phylogenetic tree** | | | | | |
| --- | --- | --- | --- | --- | --- |
| *psbA* | *psbD* | *accD* | *rps12* | *rpl22* | *ndhA* |
| *matK* | *psbC* | *psaI* | *clpP* | *rps19* | *ndhH* |
| *psbK* | *psbZ* | *cemA* | *psbT* | *rpl2* | *rps15* |
| *psbI* | *rps14* | *petA* | *psbN* | *rpl23* | *ycf1* |
| *atpA* | *psaB* | *psbJ* | *psbH* | *ycf2* |  |
| *atpF* | *psaA* | *psbL* | *petB* | *ndhB* |  |
| *atpH* | *ycf3* | *psbF* | *petD* | *rps7* |  |
| *atpI* | *rps4* | *psbE* | *rpoA* | *ndhF* |  |
| *rps2* | *ndhJ* | *petL* | *rps11* | *ccsA* |  |
| *rpoC2* | *ndhK* | *petG* | *rpl36* | *ndhD* |  |
| *rpoC1* | *ndhC* | *psaJ* | *rps8* | *psaC* |  |
| *rpoB* | *atpE* | *rpl33* | *rpl14* | *ndhE* |  |
| *petN* | *atpB* | *rps18* | *rpl16* | *ndhG* |  |
| *psbM* | *rbcL* | *rpl20* | *rps3* | *ndhI* |  |

**Table S10.** Comparison of polymorphic sites (74 CDS used in phylogenomics tree construction) between *G. mangostana* var Mesta/Manggis versus other *Garcinia* species.

| **Species** | **Number of sites** | **Sites with alignment gaps or missing data** | **Invariable sites** | **Variable sites** |
| --- | --- | --- | --- | --- |
| *G. anomala* | 66,117 | 123 (0.19%) | 65,736 (99.42%) | 258 (0.39%) |
| *G. gummi-gutta* | 66,153 | 132 (0.20%) | 65,728 (99.36%) | 293 (0.44%) |
| *G. mangostana* var *Thailand* | 66,144 | 243 (0.37%) | 65,342 (98.79%) | 559 (0.85%) |
| *G. oblongifolia* | 66,036 | 180 (0.27%) | 65,588 (99.32%) | 448 (0.68%) |
| *G. paucinervis* | 66,147 | 177 (0.27%) | 65,325 (98.76%) | 645 (0.98%) |
| *G. pedunculata* | 66,114 | 123 (0.19%) | 65,414 (98.94%) | 577 (0.87%) |

*Parentheses indicate the percentage of gaps/invariable/variable sites

**Table S11.** List of species used for phylogenetic tree construction using *ITS* gene.

| **Species** | **Locality/origin** | **GenBank/DDBJ* no.** |
| --- | --- | --- |
| *G. mangostana* SAM | South America | AJ 509214 |
| *G. mangostana* PM1 | Peninsular Malaysia | AF 367215 |
| *G. mangostana* PM2 | Peninsular Malaysia | AB 110808 |
| *G. mangostana* PM3 | Peninsular Malaysia | AB856024* |
| *G. mangostana* MASTA | Peninsular Malaysia | AB856025* |
| *G. mangostana* JAV | Java Island | AB 110807 |
| *G. mangostana* TH1 | Thailand | AB110809 |
| *G. mangostana* TH2 | Thailand | AB110810 |
| *G. mangostana* TH3 | Thailand | AB110811 |
| *G. mangostana* LAO | Laos | AB856023* |
| *G. malaccensis* MY1 | East coast, Peninsular Malaysia | AB856027* |
| *G. malaccensis* MY2 | East coast, Peninsular Malaysia | AB856028* |
| *G. malaccensis* MY3 | East coast, Peninsular Malaysia | AB856029* |
| *G. malaccensis* MY4 | North East, Peninsular Malaysia | AB856026* |
| *G. malaccensis* MY5 | South, Peninsular Malaysia | AB856030* |
| *G. malaccensis* SUM1 | Sumatra, Indonesia | AB 110805 |
| *G. malaccensis* SUM2 | Sumatra, Indonesia | AB 110806 |
| *G. malaccensis* SBH | Sabah, Malaysia | AB856031* |
| *G. penangiana* 1 | West Peninsular Malaysia | AF 367226 |
| *G. penangiana* 2 | East Peninsular Malaysia | AB856033* |
| *G. diospyrifolia* | West, Peninsular Malaysia | AF 367227 |
| *G. celebica* | Peninsular Malaysia | AB856032* |
| *G. atroviridis* | Peninsular Malaysia | AF 367211 |

Asterisk denotes sequences that were submitted to DDBJ

Without asterisk sequences refers to GenBank accessions

**Table S11 (continue).** List of species used for phylogenetic tree construction using *ITS* gene.

| **Species** | **Locality/origin** | **GenBank** |
| --- | --- | --- |
| *G. paucinervis* CH1 | China | KX421323 |
| *G. paucinervis* CH2 | China | KX421324 |
| *G. paucinervis* CH3 | China | KX421325 |
| *G. oblongifolia* | China | KX421326 |
| *G. pedunculata* InD1 | Western Ghats, India | KP318343 |
| *G. pedunculata* InD2 | Western Ghats, India | KP318344 |
| *G. pedunculata* InD3 | Western Ghats, India | KP318345 |
| *G. pedunculata* InD4 | Western Ghats, India | KP318346 |
| *G. oblongifolia* CH1 | China | KX421327 |
| *G. oblongifolia* CH2 | China | KX421328 |
| *G. gummi-gutta* InD1 | India | KY659403 |
| *G. gummi-gutta* InD2 | India | KY659404 |
| *G. gummi-gutta* InD3 | India | KX765284 |
| *Garcinia celebica* BBI1 | Indonesia: Bogor Botanic Garden | LC010524 |
| *Garcinia celebica* BBI2 | Indonesia: Bogor Botanic Garden | LC010525 |
| *Garcinia celebica* BBI3 | Indonesia: Bogor Botanic Garden | LC010527 |
| *Garcinia hombroniana* BBI1 | Indonesia: Bogor Botanic Garden | LC010528 |
| *Garcinia hombroniana* BBI2 | Indonesia: Bogor Botanic Garden | LC010529 |
| *Garcinia hombroniana* BBI3 | Indonesia: Bogor Botanic Garden | LC010530 |
| *G. mangostana* var Manggis UKM | Universiti Kebangsaan Malaysia, Bangi | OK576274 |
| *G. mangostana* var Mesta UKM | Universiti Kebangsaan Malaysia, Bangi | OK576276 |

**Table S12.** Summary of different methods used for Manggis plastome assembly.

| **Software** | | **GetOrganelle** | | **Platanus** |
| --- | --- | --- | --- | --- |
| **Assembly Method** | | **Reference-guided assembly** | ***De novo* assembly** | ***De novo* assembly** |
| Structure | | Complete circular plastome | Linear circular contig with manual curation needed | 5 scaffolds |
| No. of plastome related contig/scaffolds | | 1 | 1 | 5 |
| Contig/Scaffolds name | | *G. mangostana* var Manggis ref Mesta | *G. mangostana* var Manggis no ref | scaffold114, scaffold46654, scaffold418, scaffold46935, scaffold47072 |
| Contig/scaffolds size | | 156,581 bp | 156,663 bp | Total = 128,569 bp  scaffold114 = 85,517 bp  scaffold46654 = 22,228 bp  scaffold418 = 17,237 bp  scaffold46935 = 2,522 bp  scaffold47072 = 1,065 bp |
| Differences detected between different methods | Indel  Position*:  between 51,105-51106 | Position: 51,100-51,123  TTCGAAAAAAAAAAAAAAAAAAAT | Position: 22-45  TTCGAAAAAAAAAAAAAAAAAAAT  or  Position: 156,603-156,627  TTCGAA**T**AAAAAAAAAAAAAAAAAT | Position at scaffold114:  51,224-51,230  TTCGAA**T** |
|  | SNP  Position*:  51,136 | Position: 51,123-51,136  TCATAAAAATAAA**G** | Position: 45-58  TCATAAAAATAAA**G**  or  Position: 156,627-156,640  TCATAAAAATAAA**A** | Position at scaffold114:  51,266-51,279  TCATAAAAATAAA**G** |

* Position at plastome of *G. mangostana* var Manggis ref Mesta; Red font: Indel or SNP

**Table S13.** CDS length comparison of *Garcinia* species before and after adjustment.

| **Gene name** | ***G. anomala*** | ***G. gummi-gutta*** | ***G. mangostana*** | | ***G. oblongifolia*** | ***G. pedunculata*** | ***G. paucinervis*** |
| --- | --- | --- | --- | --- | --- | --- | --- |
|  |  |  | **Mesta/Manggis** | **Thailand** |  |  |  |
| *rps16* exon1 | 40 | 40 | 40 | 40 | 40 | 40 | 40 |
| *rps16* exon2 | 179 | 179 | 179 | 179 | 89/110 | 62 | 224 |
| *atpF* exon 1 | 145 | 145 | 145 | 145 | 145 | 145 | 144/145 |
| *atpF* exon 2 | 398 | 398 | 398 | 398 | 398 | 398 | 399/398 |
| *rpoC1* exon 1 | 432 | 432 | 432 | 432 | 432 | 432 | 430/432 |
| *rpoC1* exon 2 | 1632 | 1632 | 1635 | 1632 | 1632 | 1632 | 1634/1632 |
| *psbM* | 105 | 105 | 105 | 105 | 105 | 159/105 | 105 |
| *ndhK* | 678 | 678 | 678 | 678 | 678 | 762/678 | 678 |
| *cemA* | 687/717 | 747 | 714 | 690/726 | 750 | 690/726 | 726 |
| *clpP* exon1 | 71 | 69/71 | 71 | 69/71 | 69/71 | 71 | 69/71 |
| *clpP* exon2 | 292 | 291/292 | 292 | 291/292 | 291/292 | 292 | 291/292 |
| *clpP* exon3 | 228 | 228 | 228 | 228 | 228 | 228 | 228 |
| *petD* exon1 | 8 | 7/8 | 8 | 7/8 | 7/8 | 8 | 7/8 |
| *petD* exon2 | 526 | 524/526 | 526 | 494/496 | 524/526 | 526 | 524/526 |
| *rps19* | 150/279 | 279 | 279 | 228 | 279 | 279 | 279 |

CDS length before/after adjustment

**Table S13 (continue).** CDS length comparison of *Garcinia* species before and after adjustment.

| **Gene name** | ***G. anomala*** | ***G. gummi-gutta*** | ***G. mangostana*** | | ***G. oblongifolia*** | ***G. pedunculata*** | ***G. paucinervis*** |
| --- | --- | --- | --- | --- | --- | --- | --- |
|  |  |  | **Mesta/Manggis** | **Thailand** |  |  |  |
| *rps12** exon1 | 114 | 114 | 114 | 114 | 114 | 114 | 114 |
| *rps12** exon2 | 232 | 232 | 232 | 232 | 232 | 232 | 233/232 |
| *rps12** exon3 | 26 | 26 | 26 | 26 | 26 | 26 | 25/26 |
| *ndhD* | 1365/1503 | 1503 | 1503 | 1503 | 1503 | 1557/1503 | 1503 |
| *ndhE* | 306 | 306 | 306 | 306 | 306 | 303/306 | 306 |
| *ndhA* exon1 | 566/561 | 562 | 562 | 562 | 562 | 567/561 | 561 |
| *ndhA* exon1 | 535/534 | 533 | 533 | 533 | 533 | 534 | 534 |

CDS length before/after adjustment

*The *rps12* gene is a trans-spliced gene with the 5′ end located in the LSC region and the duplicated 3′ ends in the IR region
